# Identification of *Slit3* as a locus affecting nicotine preference in zebrafish and human smoking behaviour

**DOI:** 10.1101/453928

**Authors:** Judit García-González, Alistair J. Brock, Matthew O. Parker, Riva Riley, David Jolliffe, Ari Sudwarts, Muy-Teck Teh, Elisabeth M. Busch-Nentwich, Derek L. Stemple, Adrian R. Martineau, Jaakko Kaprio, Teemu Palviainen, Valerie Kuan, Robert T. Walton, Caroline H. Brennan

## Abstract

To facilitate smoking genetics research we determined whether a screen of mutagenized zebrafish for nicotine preference could predict loci affecting smoking behaviour. Of 30 ENU mutagenized families screened, two showed increased or decreased nicotine preference. Out of 25 inactivating mutations in the families, one in the *slit3* gene segregated with increased nicotine preference in heterozygous individuals. Focussed SNP analysis of the human *SLIT3* locus in cohorts from UK (n=863) and Finland (n=1715) identified two variants that predict cigarette consumption and likelihood of cessation. Characterisation of *slit3* mutant larvae and adult fish revealed decreased sensitivity to the dopaminergic and serotonergic antagonist amisulpride, known to affect startle reflex that is correlated with addiction in humans, and increased *htr1aa* mRNA expression in mutant larvae. No effect on neuronal pathfinding was detected. These findings reveal a role for SLIT3 in development of pathways affecting responses to nicotine in zebrafish and smoking in humans.

## INTRODUCTION

Tobacco smoking is the leading preventable cause of death worldwide placing a heavy social and financial burden on society (1–3). It is well established that aspects of smoking behaviour have a strong genetic component (4–7). However, identifying causal genetic factors and exploring the mechanisms by which they act is challenging in human studies: the field has been characterized by small effect sizes and lack of replication such that there are remarkably few genes and loci that can be confidently linked to smoking. The strongest evidence for causal effects is for functional variants in *CHRNA5* (8) and *CYP2A6* (4), affecting amount smoked and nicotine metabolism, respectively. Recent large studies have identified numerous new association loci, but their significance is yet to be biologically characterised (6, 7).

As approaches to identify genetic risk are difficult in humans, research has been facilitated by studies in animal models, with a focus on genomic analysis of inbred and selectively-bred, naturally occurring genetic strains (9). This type of study produces quantitative trait loci (QTL) maps of multiple loci, each with a small impact on the phenotype. However, as with human studies, it is inherently difficult to identify relevant genes from QTL maps, as the overall phenotype cannot be predicted by individual genotypes. Mutagenesis studies in animal model systems can overcome these limitations: e.g. N-ethyl-N-nitrosourea (ENU) mutagenesis introduces thousands of point mutations into the genome with the potential to generate much stronger phenotypes than those occurring in a natural population thereby facilitating identification of causal mutations. Examination of phenotypic variation in ENU mutagenized model species could be applied to identify novel, naturally occurring variants influencing human addictive behaviour by identifying key genes and pathways affecting conserved behavioural phenotypes.

Drug-induced reinforcement of behaviour, that reflects the hedonic value of drugs of abuse including nicotine, is highly conserved in both mammalian and non-mammalian species (10–13). Conditioned place preference (CPP), where drug exposure is paired with specific environmental cues, is commonly used as a measure of drug-induced reward or reinforcement (14). ENU Mutagenesis screens for cocaine or amphetamine-induced CPP have been undertaken in zebrafish (9, 15), however, despite successful isolation of lines with altered reinforcement responses to these drugs, the causal mutations have not been identified and the predictive validity of these screens for human behaviour has not been established. Larval locomotor response to nicotine has also been used to explore nicotine response genetics (16) but the relevance of larval locomotion to addiction is somewhat less clear.

Here, we conducted a forward genetic screen of families of ENU-mutagenized zebrafish for nicotine-induced CPP. Zebrafish express homologues of all 16 members of the nicotinic acetylcholine receptor family present in mammals (17–19) with similar binding characteristics (20, 21). However, as a result of a local gene duplication event in the ray fish lineage and a teleost genome tetraploidation event zebrafish have duplicate copies of nicotinic receptor, α2, α4, α7, α9, α10, β1, β3 and β5. In addition, zebrafish have additional receptors (α8 and α11) that have been lost in humans (19). Zebrafish show robust CPP to nicotine (21–24). Nicotinic receptor partial agonists, that modulate striatal dopamine release in response to nicotine in mammalian systems, also inhibit nicotine-induced CPP in zebrafish (21). Further, on prolonged exposure to nicotine or ethanol, adult zebrafish show conserved adaptive changes in gene expression and develop dependence-related behaviours, such as persistent drug seeking despite adverse stimuli or reinstatement of drug seeking following periods of abstinence (24). These data demonstrate the existence of a conserved nicotine-responsive reward pathway and support the suitability of zebrafish to examine the genetic and molecular mechanisms underlying behavioural responses to nicotine.

To evaluate the use of a behavioural CPP screen in zebrafish to predict loci affecting human smoking behaviour we initially assessed 1) the ability of varenicline and bupropion, pharmacological agents used to treat human nicotine addiction, to reduce zebrafish nicotine-induced place preference and 2) the heritability of nicotine responses in ENU-mutagenized fish. We then screened 30 families of ENU-mutagenized fish to identify families with increased/decreased CPP for nicotine. For two families with altered CPP response, the phenotype was confirmed following independent replication with a larger number of fish. Exome sequence information was used to generate a list of coding, loss of function candidate mutations affecting the phenotype. One family with a mutation co-segregating with increased nicotine CPP was selected for further study. Firstly, the effect of the identified gene on nicotine-induced CPP was confirmed using an independent line carrying a similar predicted loss of function mutation in the same gene. We then characterized the mutation using gene expression analysis, immunohistochemical analysis of neuronal pathways and behavioural responses to acoustic startle; a response known to be modulated by serotonergic and dopaminergic signalling and, in humans, associated with vulnerability to addiction (25–27). Finally, we used focused SNP analysis of human cohorts to assess the predictive validity of findings in fish for human smoking behaviour.

In agreement with previous studies zebrafish showed a robust CPP to nicotine. Nicotine-induced CPP was abolished by varenicline and bupropion and found to be heritable in fish. The screening of ENU mutagenized families identified mutations in the *slit3* gene influencing sensitivity to rewarding effects of nicotine. *Slit3* mutant larvae and adult fish showed reduced behavioural sensitivity to amisulpride and larvae showed increased *ht1aa* receptor expression. No effect on neuronal pathfinding was detected. Analysis of the *SLIT3* locus in two independent human cohorts identified two genetic markers that predict level of cigarette consumption and likelihood of cessation. This proof of principle study demonstrates that screening of zebrafish is able to predict loci affecting complex human behavioural phenotypes and suggests a role for SLIT3 signalling in the development of dopaminergic and serotonergic pathways affecting behaviours associated with nicotine sensitivity.

## RESULTS

### Nicotine CPP in zebrafish is inhibited by varenicline and buproprion

The hedonic value of drugs of abuse, that gives rise to reinforced behaviour, is commonly assessed using either self-administration protocols or CPP. The ability of compounds used as therapeutics in humans to prevent rodent nicotine self-administration is used to support the translational relevance of nicotine-self administration in that model (28–30). As our aim was to use nicotine-CPP to predict genes affecting smoking behaviour, we assessed the ability of the nicotine therapeutics varenicline and bupropion to inhibit nicotine induced CPP in zebrafish. As seen previously (22,24,31), 10μM nicotine induced a robust 15-20% change in place preference. Consistent with previous results (21), pre-incubation in varenicline or bupropion dose-dependently inhibited the nicotine CPP response (Figure 1).

**Figure 1.**
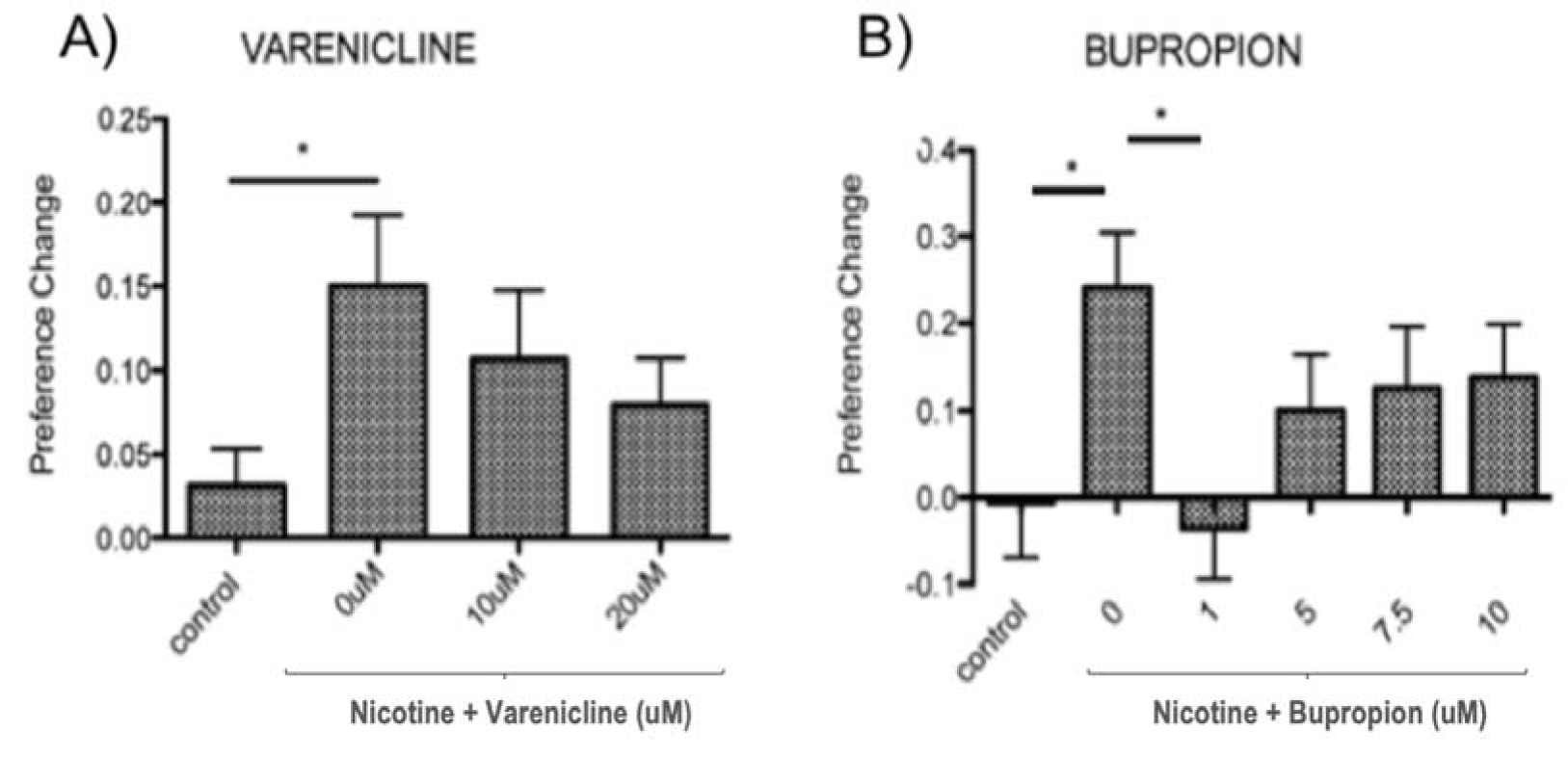
**10**μ**M nicotine induced place preference in zebrafish is sensitive to inhibition by therapeutics effective in humans.** A) varenicline (nicotine partial agonist) and B) bupropion (norepinephrine and dopamine reuptake inhibitor with nicotine antagonist properties when metabolised). Bars represent mean and error bars represent +SEM. Asterisk (*) represents significance at p<0.05.

### Nicotine CPP is heritable in zebrafish

We selected a nicotine concentration predicted to induce a minimal detectable CPP in wild types (5μM) (22, 23), to enable us to detect both increased and decreased response to nicotine in mutants. To ensure that this strategy could detect genetic factors affecting response to nicotine, we assessed the heritability of the CPP response in ENU mutagenized fish using a selective breeding approach over three generations. Figure 2A shows our assessment strategy where fish showing the highest and lowest CPP response are selected for further breeding. In the first generation, the CPP change score phenotype was normally distributed (Shapiro-Wilks p = 0.83) and there was a mean CPP change score of 0.11 to the drug paired side. CPP change scores ranged from −0.4 to 0.6.

**Figure 2.**
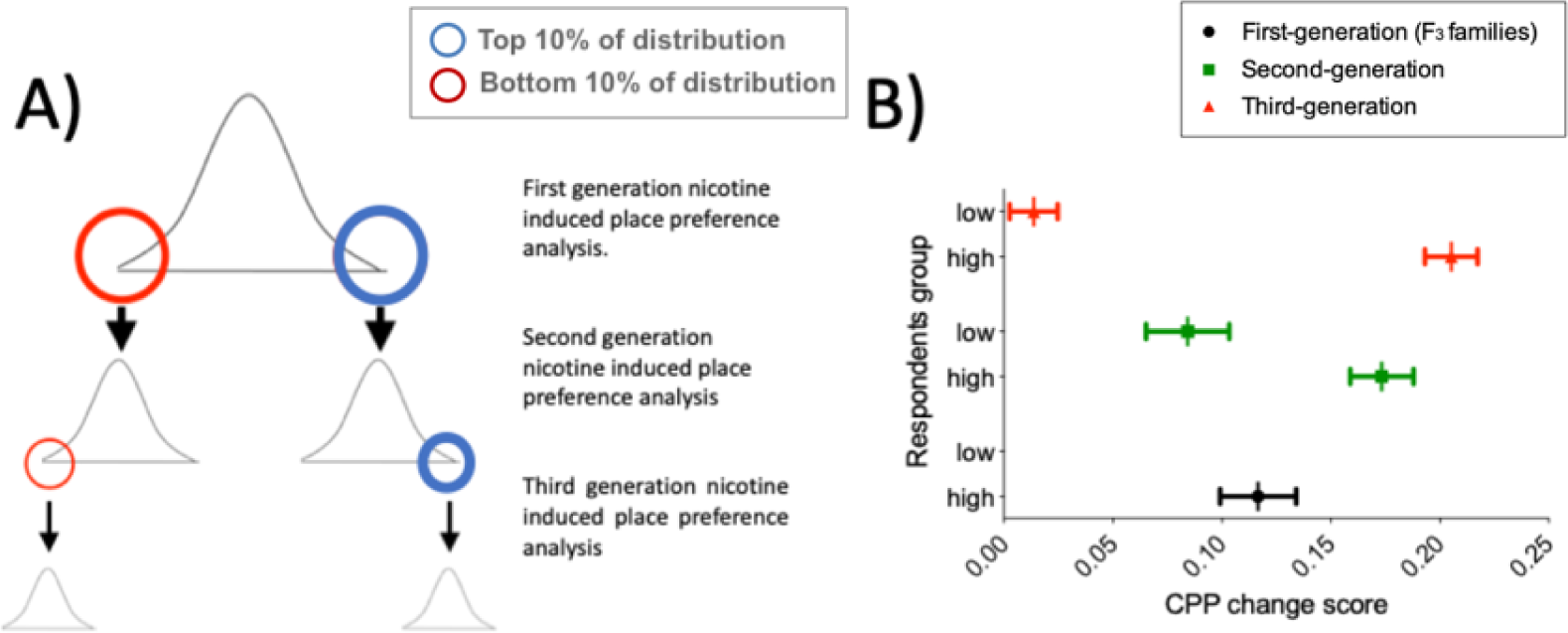
A) Breeding and selection to assess heritability of nicotine-induced place preference in ENU-mutagenized zebrafish. To test whether nicotine preference is heritable, fish in the upper (blue circle) and lower (red circle) 10% of the change in preference distribution curve were inbred and screened for CPP (Second generation CPP assay). A similar approach was used for the third generation CPP assay. **B) CPP for nicotine is heritable**. Mean preference change is increasingly distinct for the second and third generation CPP analysis. Plot represents mean and ±SEM. First generation (corresponding to the F_3_ families used for the screen) (n=120): mean=0.11; SD=0.17. Second generation: Offspring of fish from *upper* 10% of the first generation screen (n=92): mean=0.17; SD=0.14. Offspring of fish from *lower* 10% of the first generation screen (n=64): mean=0.08; SD=0.15. Third generation. Offspring of fish from *upper* 10% of the second generation screen (n=69): mean=0.21; SD=0.10. Offspring of fish from *lower* 10% of the second generation screen (n=67): mean=0.01; SD=0.09.

An increasing difference in nicotine preference between offspring of fish from the upper vs lower extremes of the distribution (Shift of Cohen’s *d* = 0.89 in Second generation CPP to *d* = 1.64 in Third generation CPP) indicates that nicotine CPP behaviour is heritable in zebrafish (Figure 2B), and that our CPP strategy is able to identify heritable differences in both extremes of the distribution. Phenotypes in the second and third generation screen may result from selecting for multiple co-segregating mutations that strengthened the phenotypes, or from selecting against other contrary mutations that weaken effects.

### Identification of *slit3* mutations affecting nicotine place preference in zebrafish

Out of 30 families screened, individuals from nine families were in the top 5% of the change in preference distribution, and individuals from five families in the bottom 5%. To identify candidate mutations affecting nicotine preference in fish, we focussed on families where all individuals included in the screen clustered at one or other extreme of the distribution curve. Two families (called AJBQM1 and AJBQM2 after the researcher who conducted the screen), which clustered at the top (AJBQM1) and bottom (AJBQM2) of the nicotine preference distribution, were selected for further study. We first assessed nicotine CPP in the remaining siblings not initially included in the screen. As shown in Figure 3, the phenotypes were conserved when remaining siblings were assessed. Exome sequencing of fish (32) used to generate AJBQM1 and AJBQM2 identified 25 nonsense and essential splice site mutations. We genotyped fish at these 25 loci and determined the co-segregation with nicotine preference.

**Figure 3.**
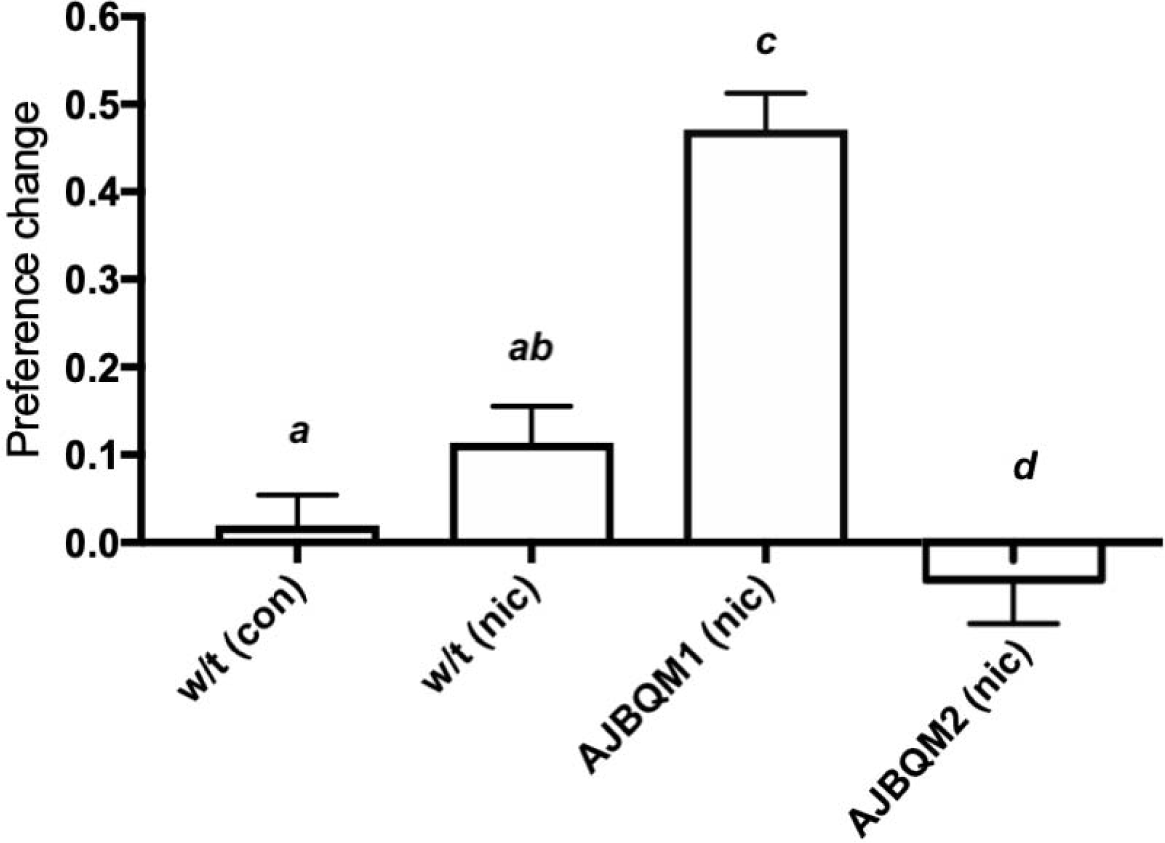
AJBQM1 and AJBQM2 families show increased and decreased nicotine place preference. AJBQM1 and AJBQM2 siblings, not included in the screen (n=10 for AJBQM1; n=14 for AJBQM2), AJBQM1 significantly differed from the parental strain, Tupfel longfin (TLF) wild type (w/t) saline control (n=17) and wild type nicotine exposed fish (n=7). AJBQM2 differed from wild type nicotine exposed fish but not wild type saline controls. Different superscript letters indicate significant difference between groups (p<0.05), same superscript letters indicate no significant differences between groups. Bars indicate Mean ±SEM.

Of the 25 coding, predicted loss of function mutations in AJBQM1 and AJBQM2 (Listed in Supplementary Table 1), only *slit3^sa1569/+^* (exon 7 splice acceptor site disruption at amino acid position 176), segregated with nicotine preference (Figure 4A & Supplementary Table 5A). None of the coding, predicted loss of function mutations in AJBQM2 segregated with nicotine preference and this line was not examined further (Supplementary Table 5B).

**Figure 4.**
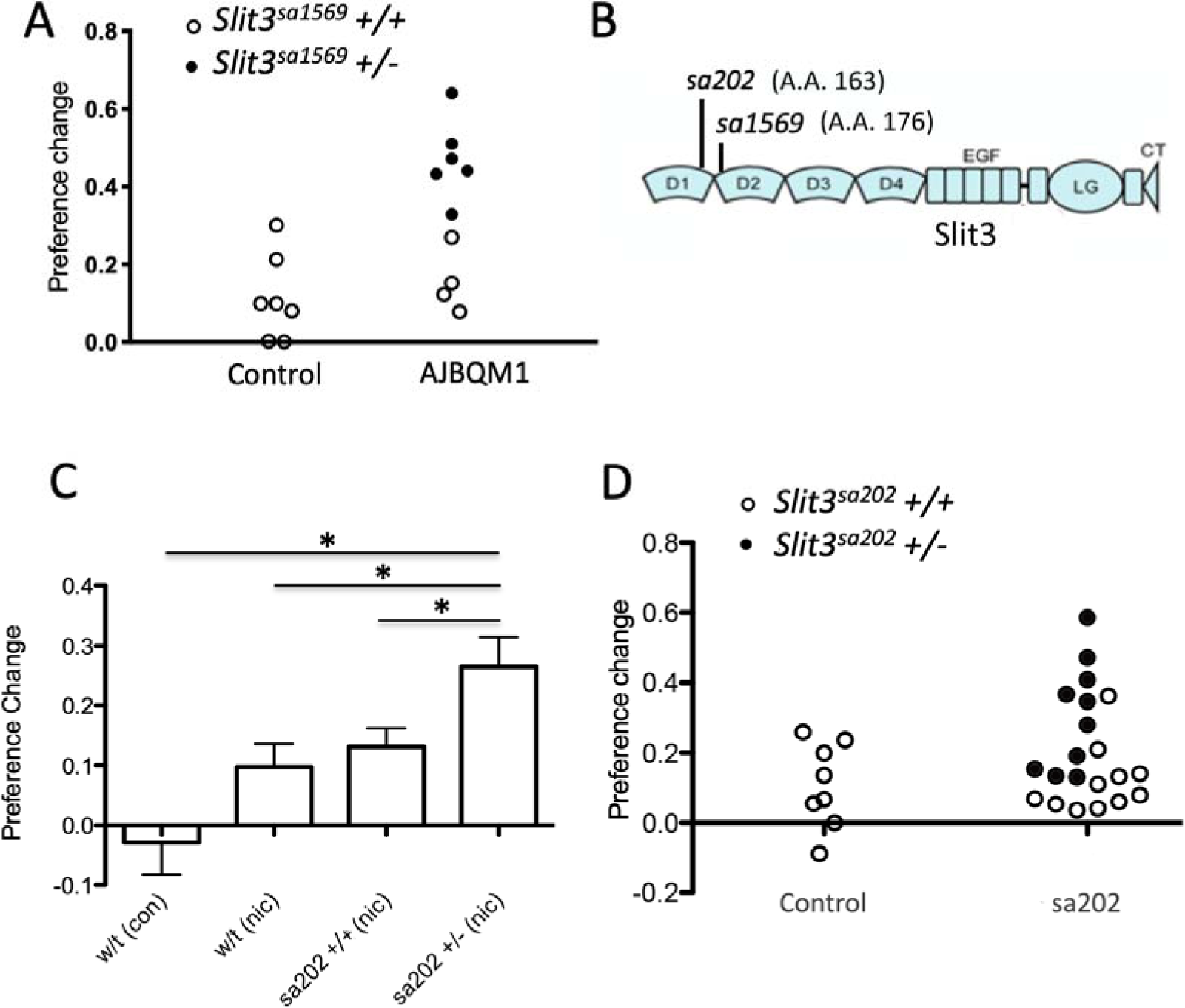
A. Segregation of *slit3^sa1569^* mutation with nicotine seeking. CPP change scores for individual un-mutagenized TLF wild type fish (n=7) and AJBQM1 fish (n=10). Following CPP analysis, fish were genotyped for 25 loss of function mutations contained within the family. Black dots indicate *slit3^sa1569/+^* heterozygous mutant fish. White dots indicate *slit3^sa1569+/+^* fish. Heterozygosity for *slit3****^sa1569^*** segregates with increased nicotine seeking behaviour. **B. Position of ENU-induced mutations in zebrafish Slit3 protein.** *slit3^sa1569^* (A>G transition) disrupts a splice site in intron 7 affecting translation at amino acid 176. *slit3^sa202^* (G>T transversion) introduces a stop codon at amino acid 163. Both mutations truncate the protein before the leucine rich repeat domain 2 (D2), which interacts with membrane bound ROBO during SLIT-ROBO signalling. **C: Nicotine preference of *slit3^sa202^* line.** *slit3^sa202/+^* fish (n=18) show increased nicotine preference compared to wild type TLF controls (n=8) (p = 0.001) and wild type siblings *slit3^+/+^* (n=14) (p<0.05). Bars indicate mean +SEM. **D: Segregation of *slit3^sa202^* allele with nicotine seeking**. CPP change scores for individual un-mutagenised TLF wild type parent strain fish (n=8) and *slit3^sa202^* fish (n=21). Black dots indicate *slit3*^sa202/+^ heterozygous mutant fish, white dots indicate *slit3^sa202+/+^* fish. Mutations in *slit3^sa202^* co-segregate with nicotine preference. Heterozygous *slit3^+/sa202^* present increased place preference compared to *slit3^sa202+/+^* siblings (n=11).

To confirm that loss of *slit3* function was related to nicotine seeking behaviour we obtained an independent family of fish, *slit3^sa202^*, carrying a G>T transversion producing a premature stop codon at amino acid position 163 in the SLIT3 protein from the Sanger Institute. Although not as marked as in AJBQM1 mutants (hereafter called *slit3^sa1569^*), heterozygous *slit3^sa202^* fish showed enhanced nicotine CPP (p=0.03) compared to wild type siblings (Figure 4C & 4D). The *slit3^sa1569^* allele affects slicing and *slit3^sa202^* introduces a premature stop codon. Both alleles reside before the second leucine rich repeat (LRR) domain in the encoded protein (Figure 4B), which is essential for interaction with ROBO receptor proteins (33).

### Characterisation of *slit3^sa1569^* mutants

SLIT3 is a member of a family of proteins with established axon guidance properties and previously suggested to be involved in dopaminergic and serotonergic pathfinding (34). Therefore, we performed immunostaining in three-day-old zebrafish larvae and examined the number of cell bodies and axonal projections of serotonergic (5-HT) and catecholaminergic neurons in the brain using anti-5-HT and anti-tyrosine hydroxylase (TH) antibodies (Figure 5).

**Figure 5:**
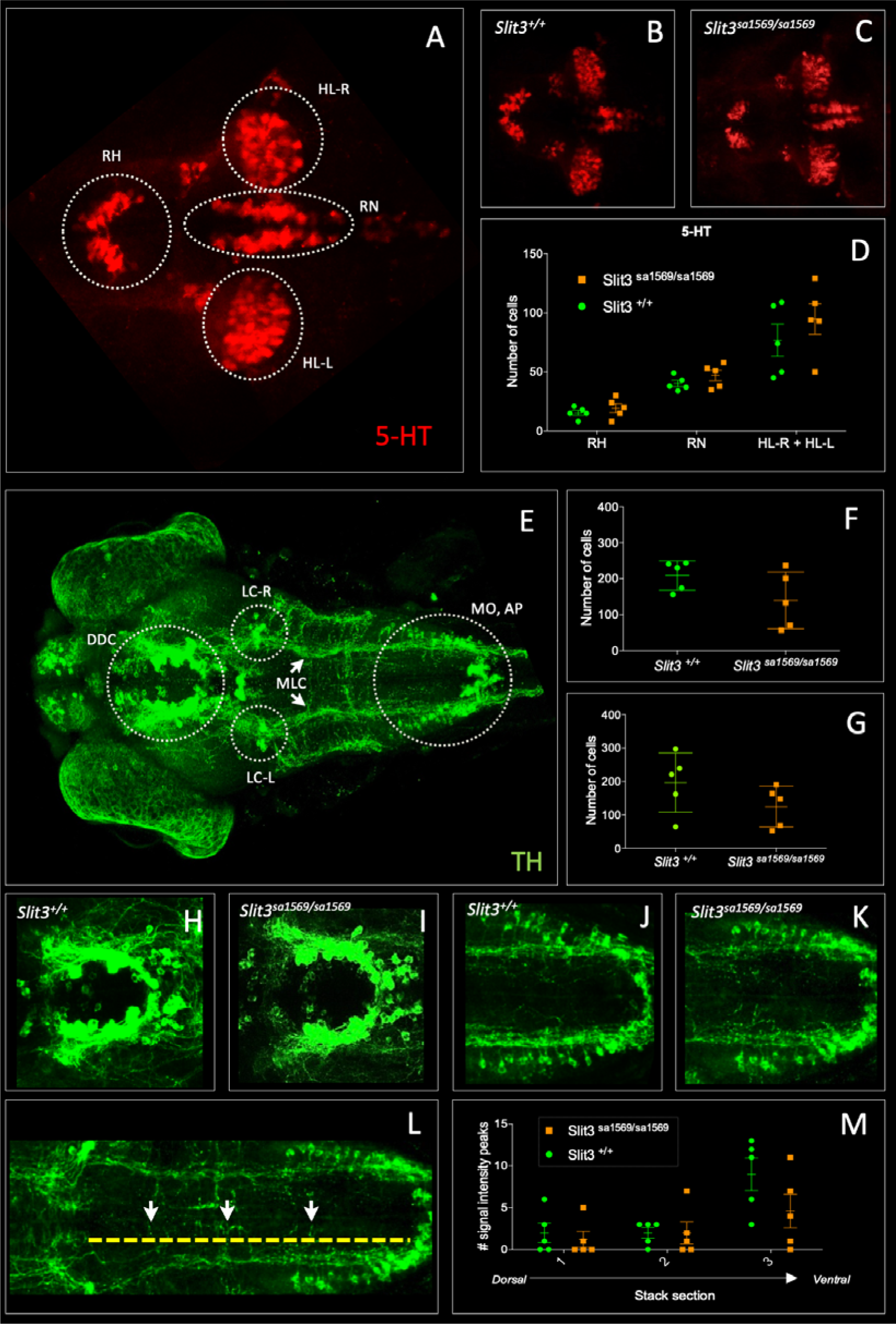
Fluorescent immunohistochemistry in three-day old wild type ***slit3^sa1569+/+^* and homozygous mutant *slit3^sa1569/sa1569^***. A-D) anti-5-HT, E-M) anti TH: A) 5-HT-labelled neurons in wild type zebrafish brain. Circles indicate regions used for quantification of cell number in rostral hypothalamus (RH), inferior hypothalamic lobes (HL-R, HL-L) and raphe nucleus (RN). B) anti-5-HT labelled cells in *slit3* wild type brain, C) anti-5-HT labelled cells in *slit3^sa1569^* homozygous mutant brain. D) Quantification of anti-5HT labelled cell number in wild type and *slit3* mutant brains No significant differences were observed between wild type and *slit3* mutant larvae. E) Unprocessed maximum intensity projection of anti-TH-labelled whole mounted wild type zebrafish brain. Circles indicate areas used for quantification, or in the case of LC-R and LC-L, landmarks used as reference to determine the extension of the medial longitudinal catecholaminergic tract (MLC) used when quantifying the number of anti-TH labelled projections to the midline (panels L, M), F) Cell quantification for diencephalic dopaminergic cluster (DDC). No significant differences were observed between wild type and *slit3* mutant larvae. G) Cell quantification for medulla oblongata interfascicular zone and vagal area, and area postrema (MO, AP). No significant differences were observed between wild type and *slit3* mutant larvae. H-K) Anti-TH labelled wild types and *slit3^sa1569^*. Zoomed-in visualization of diencephalic dopaminergic cluster (H-I) and medulla oblongata interfascicular zone and vagal area (J-K). L-M) Quantification of catecholaminergic projections projecting to the midline. Examples of projections are indicated with yellow arrows. Projections were assessed from posterior to anterior using the locus coerulus and posterior extent of the raphe nucleus as landmarks (Panel L, yellow line) and from dorsal to ventral (Panel M, stacks 1-3). Figure 5-figure supplement 1 shows individual planes. n=5 samples per genotype group.

No differences between *slit3^sa1569^* mutant and wild type larvae were observed in the number of cells labelled by anti-5HT antibody in the raphe nucleus (Mean ± SEM: *slit3^+/+^*: 40.2±2.6 vs. *slit3^sa1569/sa1569^*: 47.0±4.7, p=0.23) rostral hypothalamus (*slit3^+/+^*: 15.4±2.2 vs. *slit3^sa1569/sa1569^*: 19.4±3.8, p=0.38) or inferior hypothalamic lobes (*slit3^+/+^*: 76.8±13.5 vs. *slit3^sa1569/sa1569^*: 94.8±12.9, p=0.36), nor in the number of cells labelled by anti-TH antibody in the diencephalic dopaminergic cluster (*slit3^+/+^*: 209±18 vs. *slit3^sa1569/sa1569^*: 140±35, p=0.12) and medulla oblongata interfascicular zone and vagal area (*slit3^+/+^*: 196±40 vs. *slit3^sa1569/sa1569^*: 124±27, p=0.17). Similarly, we observed no significant differences in the number of anti-TH labelled axon tracts projecting to the midline across the 3 planes examined (*slit3^+/+^*: 2.6±2.3 vs *slit3^sa1569/sa1569^*: 4.3±1, p=0.53) (Figure 5).

We also looked at the expression patterns using anti-acetylated tubulin antibody along the midline in the ventral forebrain, where *slit3* is known to be expressed (35). However, no obvious differences were observed. Staining of *slbp^ty77e/ty77e^* mutant larvae, known to have fewer neurons and axonal defects (16) were used as positive control (Figure 5-figure supplement 1).

Although we did not observe differences between wild type and *slit3^sa1569^* mutants, subtle effects on circuit formation may not have been detected by our antibody staining. To further characterise the *slit3* phenotype and explore potential functional differences in catecholamine circuitry, we examined the response and habituation to acoustic startle stimuli in wild type and mutant fish. Habituation to acoustic startle is known to involve catecholamine signalling and to be sensitive to dopaminergic/serotonergic antagonists such as amisulpride (36) and, in humans, is associated with vulnerability to addiction (25–27). Five day old larvae were subjected to 10 sound/vibration stimuli over a total of 20 seconds (2 second interval between each stimulus) in the presence of 0, 0.05 mg/L, 0.1 mg/L or 0.5 mg/L amisulpride in 0.05% dimethyl sulfoxide (DMSO). The distance travelled one second after each stimulus was recorded for each fish.

Response and habituation to the stimuli was quantified as the percentage of fish moving more than 4.6 mm, which corresponds to 85% of the mean distance travelled one second after the first startle (Figure 6A). Using this criterion – and in line with the habituation response paradigm (37) – a lower percentage of fish responded as the number of stimuli increased: 68% or wild type non drug-treated individuals responded to the first stimulus, 81% to the first and/or second stimulus, whereas 16% responded to the last stimulus (Figure 6B).

**Figure 6.**
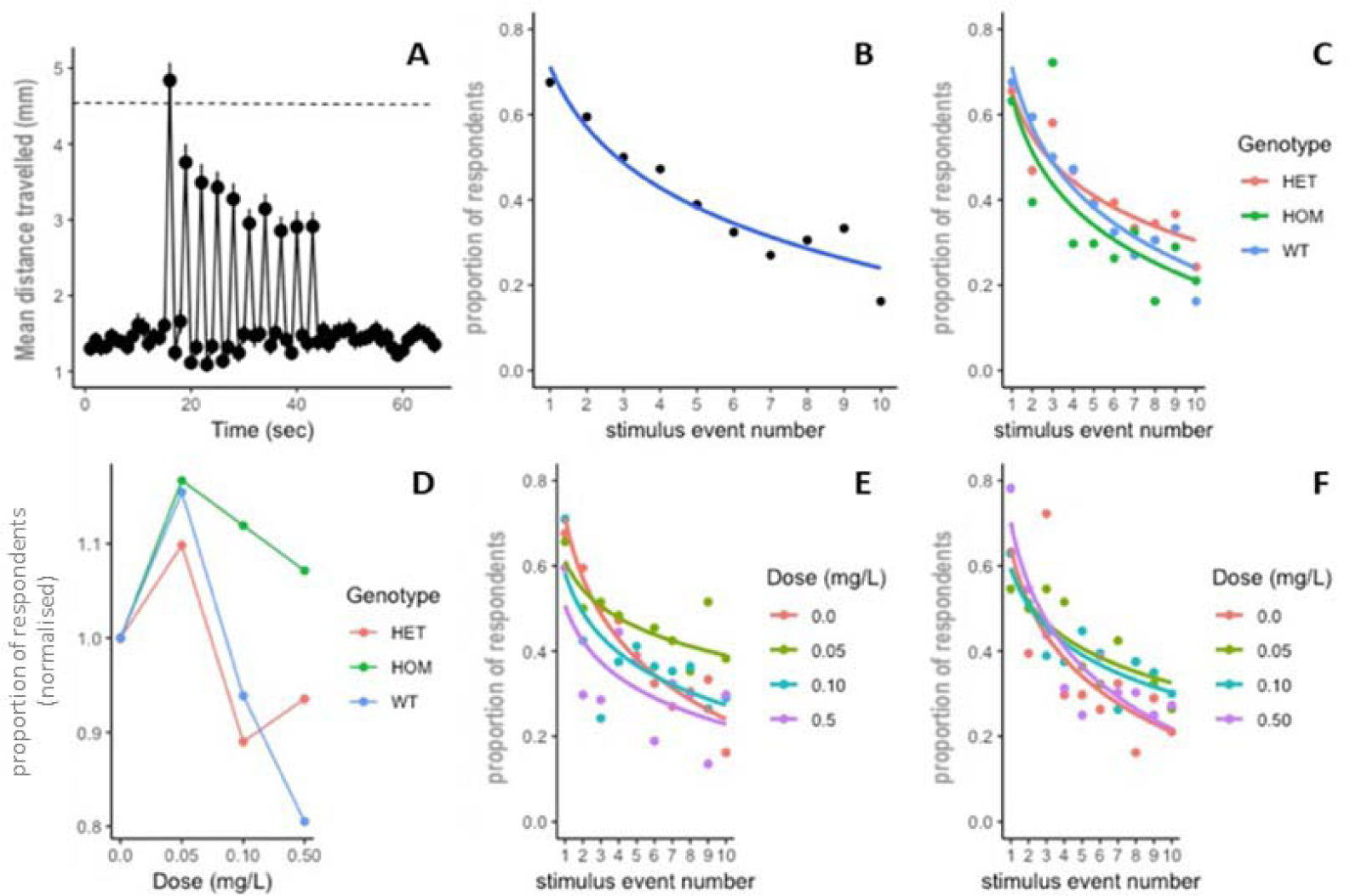
**Habituation response in the presence and absence of amisulpride. A-B: Response and habituation to 10 stimuli with two seconds interval between stimuli in wild type, drug free zebrafish**. Mean distances travelled were measured in one second time bins. Line indicates 4.6 mm, which corresponds to 85% of the population mean distance travelled one second after the first stimulus and was used to define respondents. The percentage of fish responding to the stimuli decreases with stimulus/tap number (Main effects of tap number p<0.05) 67% respond to the first tap; 16% respond to the last tap. Respondents are defined as fish moving more than 4.6 mm. **C: proportion of responders across the ten stimuli in drug free individuals from each genotype:** there was no significant effect of genotype on response across taps (p=0.340) or responsiveness (p=0.352) in drug free fish. **D: Mean percentage of responders across the ten stimuli (±SEM).** Data are stratified by *slit3^sa1569^* genotype and amisulpride dose normalised to response in absence of drug. The effect of amisulpride on habituation varies by genotype. **E/F: proportion of individuals responding in each amisulpride dose condition in wild type and homozygous mutant fish, respectively.** The interaction between amisulpride dose and stimulus event number had a significant effect on the proportion of responsive individuals in wild type individuals (p<0.05) but not homozygous mutants (p=0.160).

In drug-free conditions, there were no differences across *slit3^sa1569^* genotype groups (Figure 6C). However, when larvae were treated with amisulpride, the habituation to startle responses across taps was different across genotypes. Amisulpride caused a biphasic dose dependent effect on stimulus response in wild types such that 0.05mg/L caused an increase in responders across all 10 stimuli, and 0.5mg/L caused a decrease (Effect of amisulpride dose p<0.001). A similar pattern was observed for heterozygous *slit3^sa1569^,* but the effect of amisulpride was not significant (p=0.083).

Amisulpride dose had no significant effect on stimulus response in homozygous *slit3^sa1569^*, that showed an increase in response to low doses but were less sensitive to inhibition at high doses (Figure 6D-F). The presence of a *slit3^sa1569^* genotype by amisulpride dose interaction across taps was confirmed by a three-way interaction in the regression models (p=0.04). The interaction between dose and stimulus event number was a significant predictor of response in wild type larvae (p=0.044) and heterozygous larvae (p=0.02) but not in homozygous larvae (p=0.16).

There were no significant differences in locomotion in the 15 seconds before the first startle, in magnitude of the response to the first tap stimulus, nor in total distance moved across all tap stimuli across experimental groups (Figure 6-figure supplement 1) indicating that differences in behaviour were not confounded by differences in locomotion per se.

Adult *slit3 ^sa1569/ sa1569^* mutant zebrafish showed a qualitatively different response to inhibition of CPP by amisulpride compared to wild type siblings, consistent with a persistent difference in sensitivity to this drug. The minimal CPP induced by 5μM nicotine in wild type fish was prevented by pre-exposure to 0.5mg/L amisulpride. Nicotine-induced CPP in *slit3^sa1569^* homozygous mutants was not affected (Figure 7).

**Figure 7.**
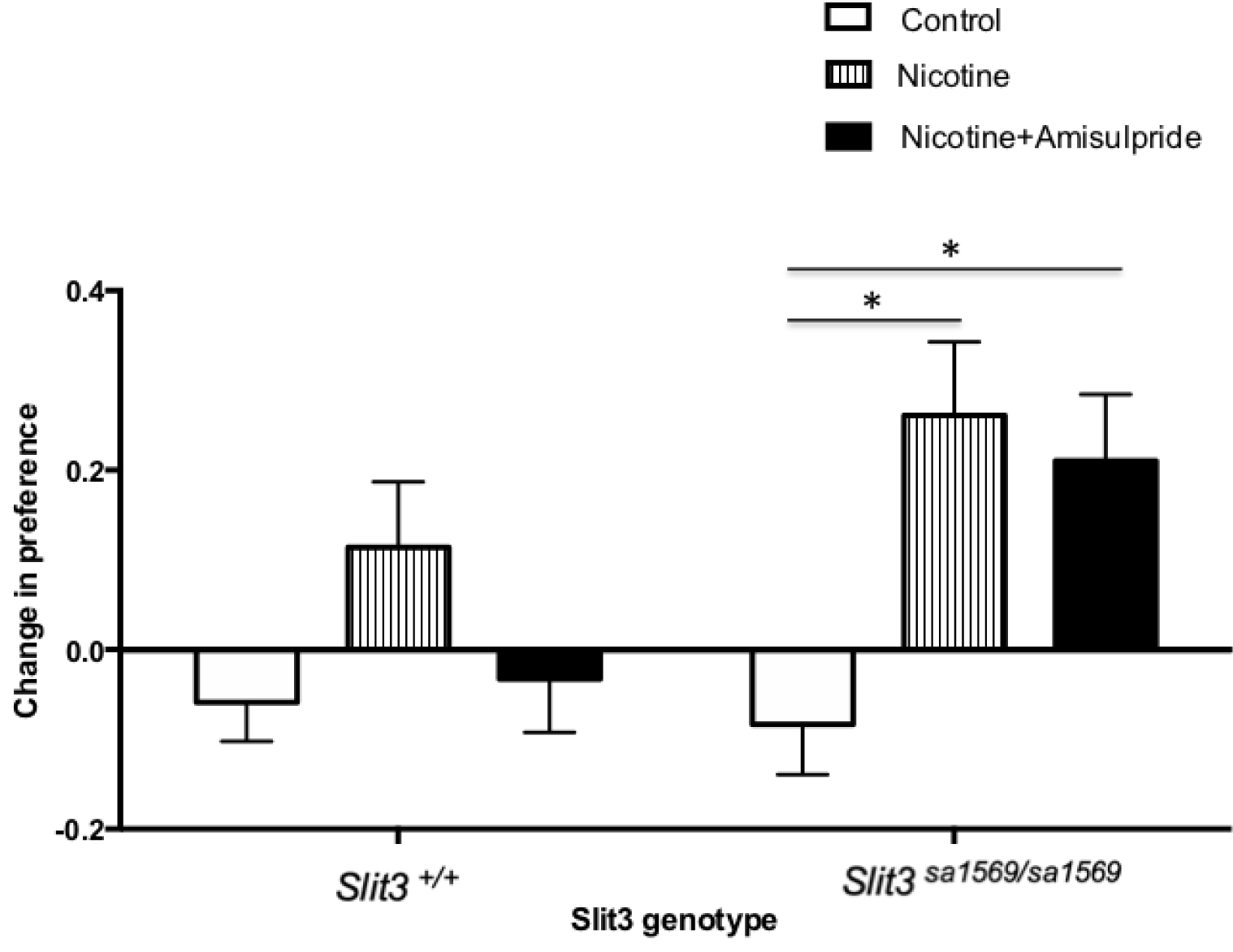
**CPP induced by 5**μM nicotine is blocked by 0.5mg/L dopamine/serotonin antagonist amisulpride in wild type *slit3^sa1569+/+^* fish but not in *slit3^sa1569^* homozygous mutants. Bars represent mean (+SEM). (n=11-14 fish per group). *Two-way ANOVA followed by post-hoc Tukey tests (p < 0.05).

As *slit3^sa1569^* homozygous mutant fish showed altered sensitivity to nicotine and amisulpride, we examined whether expression of genes previously associated with nicotine dependence was dysregulated in *slit3^sa1569^* mutant larvae using quantitative real-time PCR (qPCR). We included the nicotinic receptors and genes from the dopamine receptor family -*drd1* (40–42)(38), *drd2* (39) and *drd3* (40)-, and the dopamine transporter *dat* (41)). Genes from the adreno-receptor families (*adra1* and *adra2*) were also included due to their links with nicotine addiction and use as putative targets for smoking treatment (43–47)(42–44). Finally, expression of serotonin receptor genes was included, again due to their well-established links with nicotine addiction (45–49).

For several genes (i.e. *drd3, chrnb3, htr4, adra2b*), up-regulation of gene expression in mutant larvae showed nominal significance (Supplementary Table 8). However, only *htr1aa* ([F(2, 6)=44], p=0.0003) showed a significant difference across genotypes after correcting for multiple testing (Figure 8).

**Figure 8.**
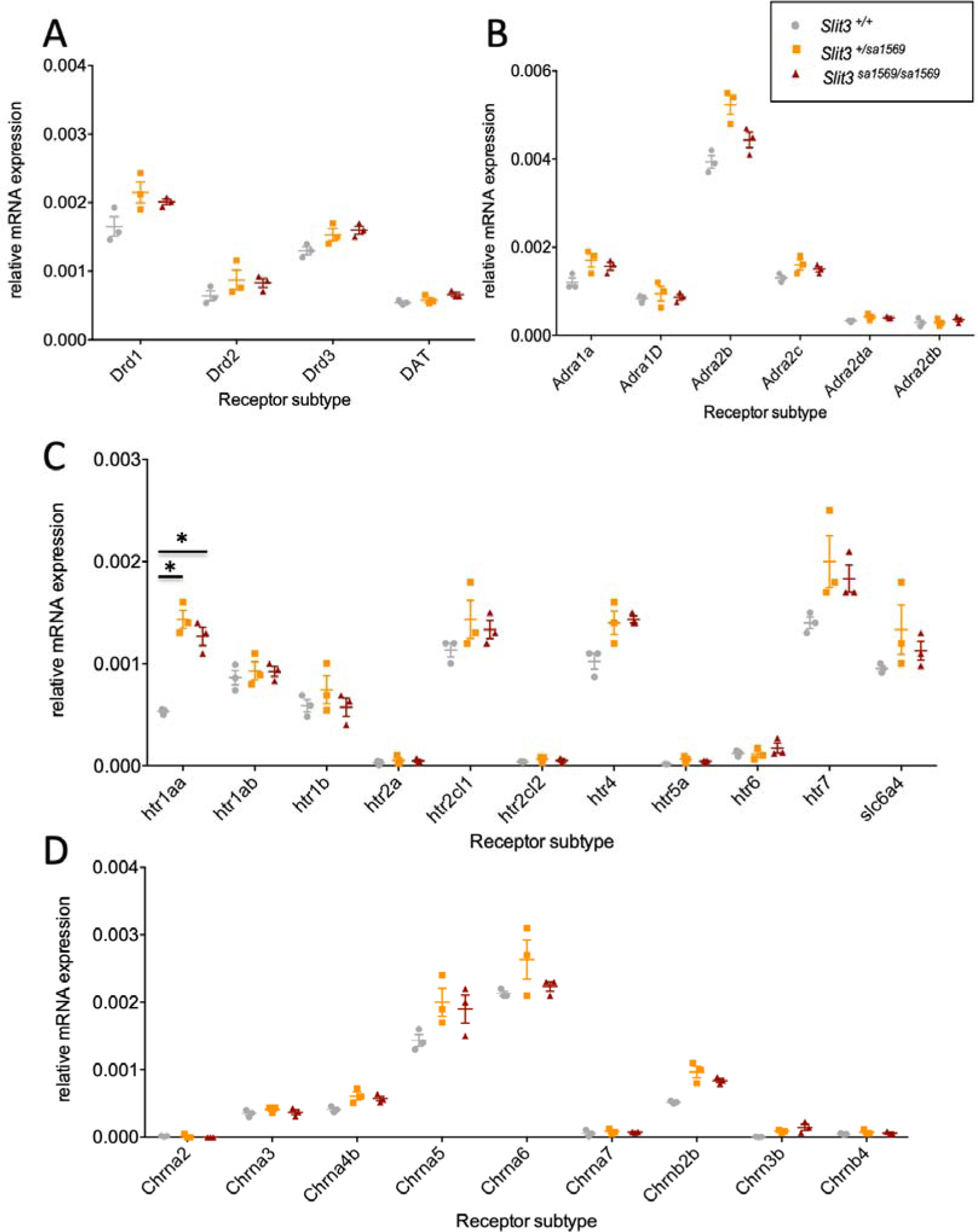
Quantitative real-time PCR analysis of five day old wild type *slit3^sa1569+/+^*, *slit3^sa1569/+^* heterozygous and *slit3^sa1569/sa1569^* homozygous mutant larvae (Total n=30, 3 samples per experimental group with n=10 embryos per sample). Only *htr1aa* ([F(2,6)=44], p=0.0003) showed a significant difference across genotypes after correcting for multiple testing. *Two-way ANOVA followed by post-hoc Tukey test (p < 0.05).

### Variations at the *SLIT3* locus predict smoking behaviour in human samples

We next examined associations between 19 single nucleotide polymorphisms (SNPs) in the human *SLIT3* gene and smoking behaviour in two London cohorts. Two SNPs, rs12654448 and rs17734503 in high linkage disequilibrium (Figure 9) were associated with level of cigarette consumption (p=0.00125 and p=0.00227). We repeated the analysis on heavy smokers: rs12654448 (p=0.0003397) and rs17734503 (p=0.0008575) were again associated with cigarette consumption together with rs11742567 (p=0.004715). The SNP rs11742567 was associated with cigarette consumption in light smokers (<20 cigarettes per day, p=0.003909)) and with quitting. Associations are reported in Table 1. No other *SLIT3* polymorphisms were associated with smoking initiation, persistent smoking or cessation (Supplementary Tables 6 & 7).

**Figure 9:**
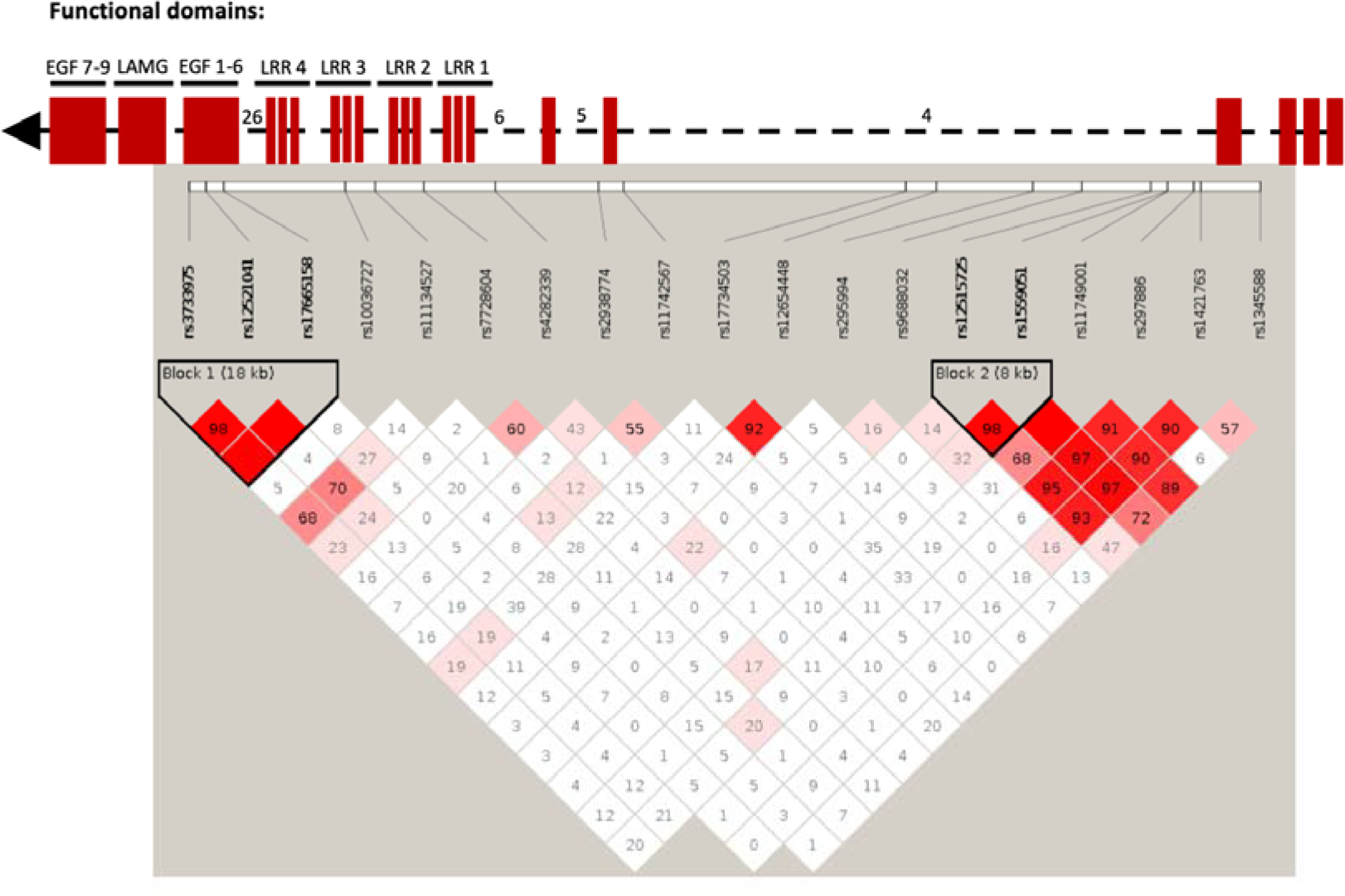
Linkage disequilibrium (LD) plot of *SLIT3* SNPs in human smoking association analysis. Numbers within each square indicate D’ values (white: D’ < 1, LOD < 2; blue: D’ = 1, LOD < 2; pink: D’ < 1, LOD ≥ 2; and bright red: D’ = 1, LOD ≥ 2). Top part of the figure shows domain organization of the SLIT protein based on ≥ the UCSC Genome Browser (http://genome.ucsc.edu/) in relation to the SNP location. LRR: leucin-rich repeats. EGF: epidermal growth factor domains. LamG; Laminin G domain. Some intron numbers were added for reference.

**Table 1.** Associations of SLIT3 SNPs with level of tobacco consumption for the London study groups (n=863). Regression coefficients, confidence intervals and p-values from linear regression of cigarettes smoked per day (CPD) on minor allele count for smokers from COPD, asthma and general cohorts, adjusted for age, sex and cohort. β coefficient represents effect of each additional minor allele. Benjamini-Hochberg cut-off at q-value 0.1 = 0.01053. **Associations of SLIT3 SNPs with tobacco consumption in a subset of heavy smokers (**≥**20 cigs/day).** Adjusted for age, sex and cohort. (q-value 0.1 = 0.01579). **Associations of SLIT3 SNPs in a subset of light smokers (<20 cigs/day).** Adjusted for age, sex and cohort (q-value 0.1 = 0.00526). **Association analysis of SLIT3 SNPs with smoking cessation.** Logistic regression of current smokers *vs* ever smokers controlling for age, sex and cohort. Odds ratio >1 indicates minor allele increases odds of persistent smoking relative to major allele. P: p-value, SE: standard error, L95: lower limit of 95% confidence interval, U95: upper limit. For all panels, associations in bold remained significant after adjustment for multiple comparisons using a Benjamini-Hochberg procedure to control false discovery rate at 10%.

|  | *Tobacco consumption* |  |  |  | *Tobacco consumption - heavy smokers (≥20 cigs/day)* |  |  |  | *Tobacco consumption - light smokers (<20 cigs/day)* |  |  |  | *Smoking cessation* |  |  |  |  |
| --- | --- | --- | --- | --- | --- | --- | --- | --- | --- | --- | --- | --- | --- | --- | --- | --- | --- |
| SNP | P value | β | SE | 95% CI | P value | β | SE | 95% CI | P value | β | SE | 95% CI | OR | SE | L95 | U95 | P value |
| rs10036727 | 0.629 | -0.388 | 0.802 | (-1.960, 1.183) | 0.448 | -0.653 | 0.860 | (-2.337, 1.032) | 0.940 | -0.051 | 0.686 | (-1.396, 1.293) | 0.947 | 0.160 | 0.693 | 1.295 | 0.734 |
| rs11134527 | 0.218 | 1.014 | 0.822 | (-0.596, 2.625) | 0.327 | -0.867 | 0.883 | (-2.599, 0.864) | 0.261 | 0.795 | 0.705 | (-0.586, 2.176) | 0.665 | 0.165 | 0.482 | 0.918 | 0.013 |
| **rs11742567** | 0.135 | -1.166 | 0.779 | (-2.691, 0.361) | **0.005** | **-2.346** | **0.825** | **(-3.962, -0.730)** | **0.004** | **1.888** | **0.644** | **(0.6258, 3.151)** | **1.586** | **0.163** | **1.153** | **2.183** | **0.005** |
| rs11749001 | 0.059 | 1.972 | 1.044 | (-0.074, 4.018) | 0.873 | 0.177 | 1.103 | (-1.985, 2.338) | 0.206 | 1.200 | 0.944 | (-0.651, 3.051) | 0.953 | 0.206 | 0.637 | 1.426 | 0.817 |
| rs12515725 | 0.592 | -0.406 | 0.756 | (-1.888, 1.076) | 0.488 | -0.565 | 0.813 | (-2.159, 1.029) | 0.278 | -0.688 | 0.631 | (-1.925, 0.550) | 1.028 | 0.151 | 0.765 | 1.381 | 0.855 |
| rs12521041 | 0.904 | -0.105 | 0.865 | (-1.801, 1.591) | 0.996 | -0.005 | 0.942 | (-1.851, 1.841) | 0.059 | -1.354 | 0.710 | (-2.746, 0.038) | 1.554 | 0.178 | 1.096 | 2.205 | 0.013 |
| **rs12654448** | **0.001** | **-4.241** | **1.307** | **(-6.803, -1.680)** | **0.0003** | **-4.830** | **1.334** | **(-7.444, -2.216)** | 0.410 | -1.034 | 1.251 | (-3.486, 1.417) | 1.625 | 0.279 | 0.941 | 2.808 | 0.082 |
| rs1345588 | 0.240 | -1.268 | 1.078 | (-3.380, 0.845) | 0.253 | -1.334 | 1.164 | (-3.616, 0.948) | 0.869 | -0.150 | 0.907 | (-1.927, 1.627) | 1.417 | 0.222 | 0.918 | 2.189 | 0.116 |
| rs1421763 | 0.272 | -0.982 | 0.894 | (-2.735, 0.770) | 0.978 | -0.027 | 0.959 | (-1.908, 1.853) | 0.162 | -1.074 | 0.764 | (-2.571, 0.424) | 0.917 | 0.176 | 0.649 | 1.294 | 0.622 |
| rs1559051 | 0.961 | -0.040 | 0.819 | (-1.644, 1.564) | 0.458 | 0.656 | 0.882 | (-1.073, 2.384) | 0.507 | 0.455 | 0.685 | (-0.880, 1.797) | 0.919 | 0.163 | 0.668 | 1.265 | 0.606 |
| rs17665158 | 0.131 | 1.338 | 0.884 | (-0.394, 3.070) | 0.236 | 1.114 | 0.939 | (-0.727, 2.955) | 0.034 | 1.620 | 0.758 | (0.1354, 3.106) | 0.723 | 0.172 | 0.516 | 1.013 | 0.060 |
| **rs17734503** | **0.002** | **-3.987** | **1.299** | **(-6.534, -1.441)** | **0.001** | **-4.458** | **1.325** | **(-7.055, -1.861)** | 0.410 | -1.034 | 1.251 | (-3.486, 1.417) | 1.616 | 0.275 | 0.942 | 2.773 | 0.081 |
| rs2938774 | 0.140 | 1.101 | 0.745 | (-0.359, 2.562) | 0.528 | 0.496 | 0.786 | (-1.044, 2.036) | 0.015 | -1.655 | 0.674 | (-2.976, -0.333) | 0.753 | 0.148 | 0.563 | 1.007 | 0.056 |
| rs295994 | 0.714 | 0.283 | 0.770 | (-1.227, 1.793) | 0.643 | 0.378 | 0.813 | (-1.215, 1.971) | 0.238 | -0.796 | 0.672 | (-2.114, 0.521) | 0.799 | 0.154 | 0.591 | 1.082 | 0.147 |
| rs297886 | 0.620 | 0.442 | 0.890 | (-1.303, 2.187) | 0.961 | -0.048 | 0.986 | (-1.979, 1.884) | 0.489 | 0.488 | 0.704 | (-0.891, 1.867) | 1.108 | 0.177 | 0.784 | 1.568 | 0.561 |
| rs3733975 | 0.909 | -0.099 | 0.860 | (-1.784, 1.587) | 0.982 | 0.022 | 0.934 | (-1.809, 1.852) | 0.059 | -1.354 | 0.710 | (-2.746, 0.038) | 1.488 | 0.176 | 1.054 | 2.101 | 0.024 |
| rs4282339 | 0.669 | -0.434 | 1.013 | (-2.419, 1.552) | 0.942 | -0.080 | 1.103 | (-2.241, 2.081) | 0.238 | -1.006 | 0.849 | (-2.670, 0.658) | 0.984 | 0.203 | 0.661 | 1.464 | 0.936 |
| rs7728604 | 0.701 | 0.286 | 0.744 | (-1.173, 1.745) | 0.321 | 0.827 | 0.832 | (-0.803, 2.457) | 0.654 | 0.262 | 0.583 | (-0.880, 1.404) | 0.935 | 0.149 | 0.698 | 1.253 | 0.653 |
| rs9688032 | 0.948 | -0.050 | 0.766 | (-1.551, 1.451) | 0.770 | 0.246 | 0.839 | (-1.398, 1.890) | 0.080 | -1.076 | 0.610 | (-2.272, 0.119) | 1.066 | 0.156 | 0.786 | 1.446 | 0.680 |

We subsequently investigated associations with more detailed smoking phenotypes in the Finnish twins cohort (50) (Table 2). Associations were observed between rs17734503 and DSM-IV nicotine dependence symptoms (p=0.0322) and age at onset of weekly smoking (p=0.00116) and between rs12654448 and age at onset of weekly smoking (p=0.00105). Associations were seen elsewhere between *SLIT3* markers and Fagerström Test for Nicotine Dependence (FTND), cigarettes smoked each day, sensation felt after smoking first cigarette and time to first cigarette in the morning. In keeping with the London studies the minor allele was associated with a lower degree of dependence and decreased cigarette consumption.

**Table 2:**
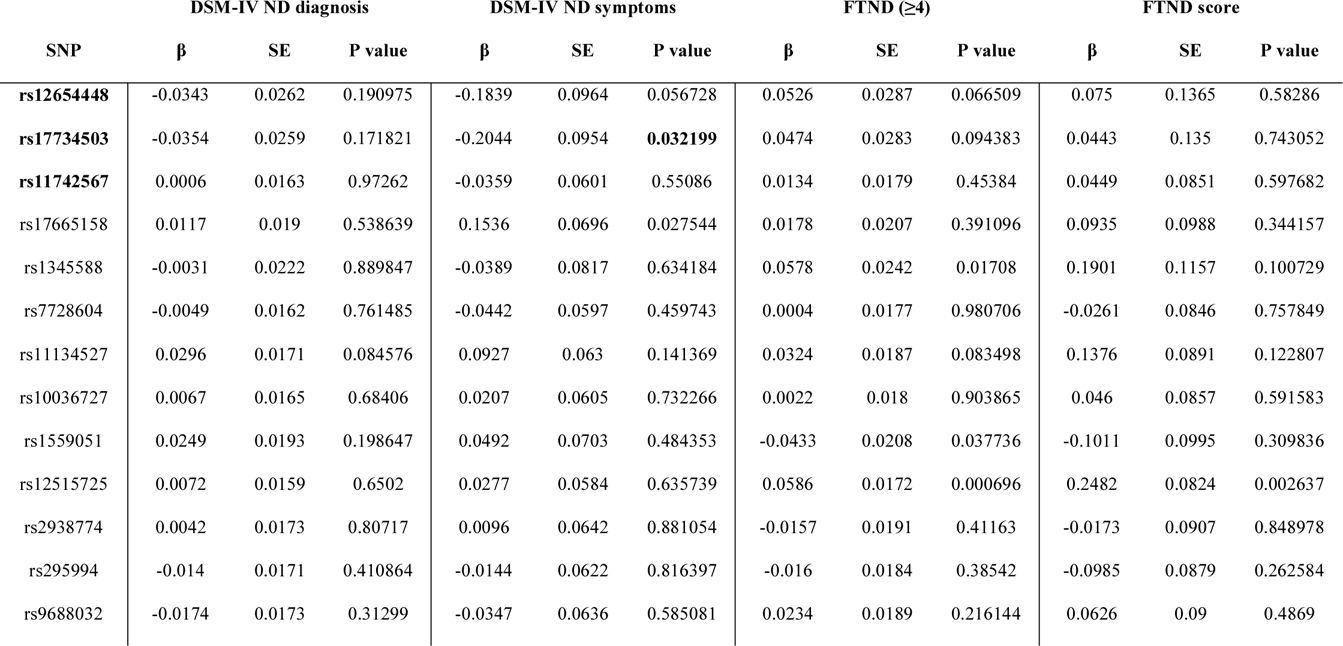
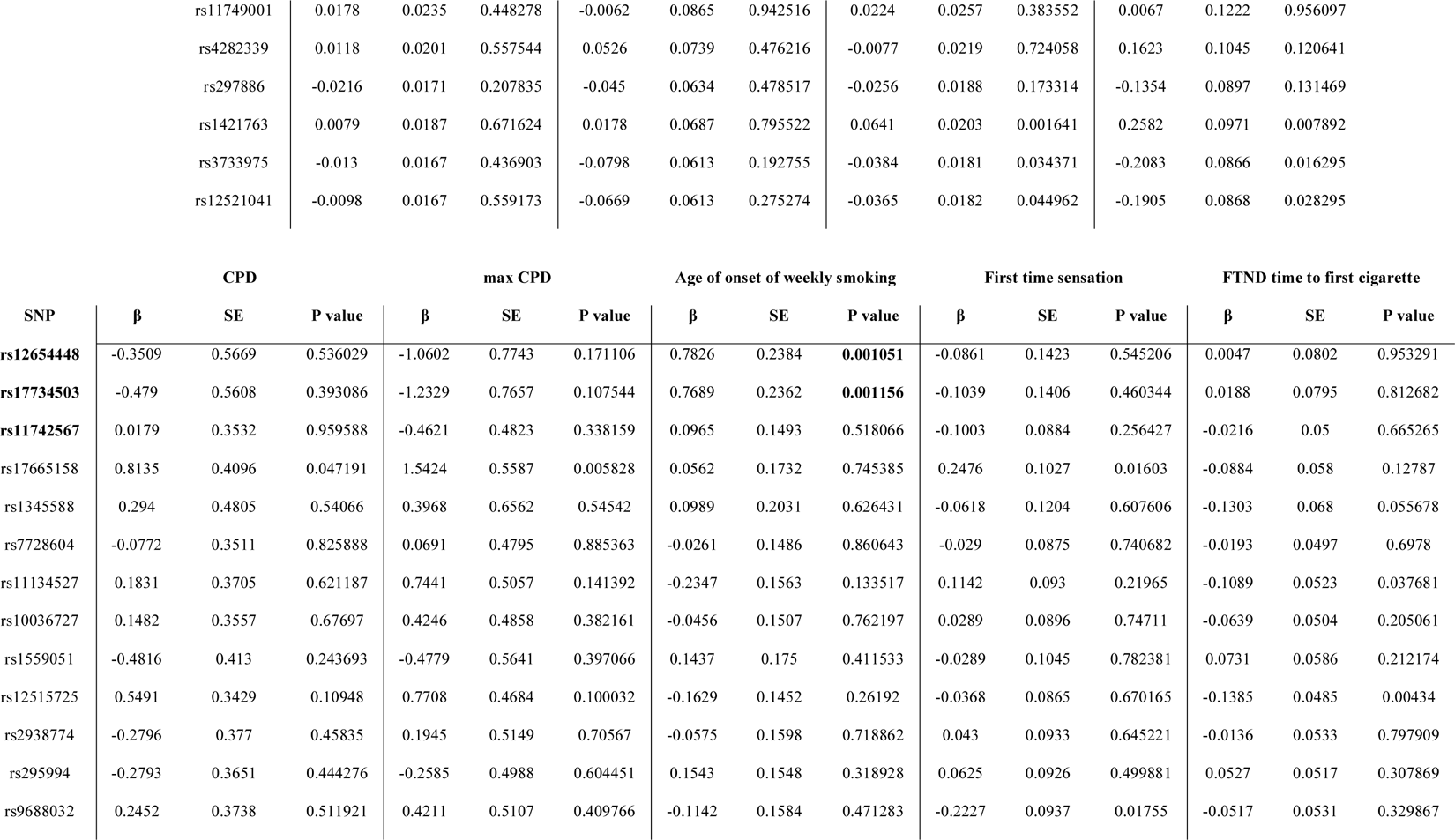
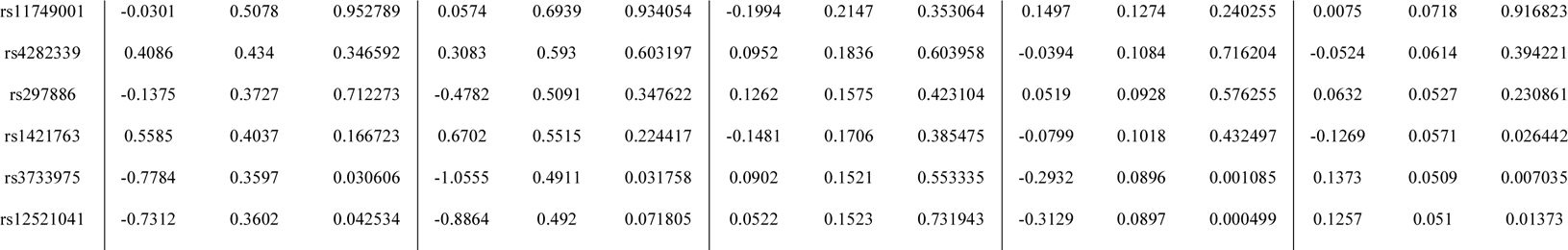
A**s**sociations **between detailed nicotine dependence phenotypes and SLIT3 genotype in a Finnish twin cohort** (n=1715). Associations of *SLIT3* SNPs with DSM-IV nicotine dependence symptoms, Fagerström scores (FTND), cigarettes smoked each day (CPD), and sensation felt after smoking first cigarette and time to first cigarette in the morning. The three SNPs that were linked to smoking behaviour in the London cohorts are shown in bold.

The SNPs rs12654448 and rs17734503 are in non-coding domains, therefore it was not possible to predict loss or gain of function of SLIT3 from the SNP location. No evidence of affecting gene expression was found as per GTEx database (https://gtexportal.org/home/).

## DISCUSSION

The aim of this study was to use forward genetic screening in zebrafish to identify loci affecting human smoking behaviour. Among 30 mutant families screened for CPP, we identified one family showing increased nicotine preference compared to wild types. Out of the 13 pre-identified loss-of-function mutations in that family, only one in the *slit3* gene co-segregated with the behaviour. We confirmed the association between *slit3* loss of function and increased nicotine preference using an independent zebrafish mutant line with a different loss of function mutation in *slit3.* Next, we established the relevance in humans by identifying two markers in *SLIT3* where the presence of the minor allele was associated with fewer cigarettes smoked each day and with smoking cessation. Studies in a separate twin cohort showed that these same alleles were associated with DSM-IV nicotine dependence symptoms and age at onset of weekly smoking. Taken together these findings suggest that zebrafish can be used to identify genes associated with smoking behaviour in humans and that variants in the *slit3* gene are linked in humans with a disruption of SLIT3 function that may affect propensity to develop tobacco dependence.

Consistent with previous findings in zebrafish and other species (21), varenicline and bupropion inhibited nicotine induced place preference in zebrafish. The pattern of inhibition differed such that varenicline (partial agonist) showed increasing inhibition at increasing concentrations, whereas inhibition by bupropion (re-uptake inhibitor) was maximal at 1μM, decreasing as concentration increased. The difference in response profile presumably reflects differences in the modes of action of these two compounds.

Our screening of ENU-mutagenized zebrafish families followed by sibling re-screen is a proof of principle study that indicates the relevance of zebrafish for human studies and emphasizes the advantage of first using a screen with a low number of individuals per family to increase efficiency. The classic three generation forward genetic screen examines phenotypes in groups of 20 or more individuals from each family (51). Logistical considerations make it difficult to apply such an approach to adult behavioural screens. Our approach increases efficiency by initially screening a small number of individuals from a large number of families and only selecting those families that occur at the extremes of the distribution for further analysis. Although in this study we were able to confirm phenotypes using a relatively small population of siblings, re-screening of a larger number would increase the power of the analysis and allow more subtle phenotypes to be identified.

One limitation of our forward genetic approach is that a large number of genes are duplicated in zebrafish due to the teleost tetraploidization; ENU-mutagenesis in one copy may not be sufficient to produce behavioural changes due to genetic compensation from the other copy of the same gene. *Slit3* is present as a single copy in the zebrafish genome which may have facilitated our ability to identify its role in responses to nicotine. However, arguably the most significant consequence of the teleost tetraploidization is the temporal and spatial specific expression of the gene duplicates. Spatial and/or temporal differences in gene expression patterns may offset concerns regarding compensation, and offer great potential to study region-specific functionality.

We identified a loss of function mutation in the zebrafish *slit3* gene associated with increased nicotine place preference and confirmed the phenotype in an independent line. SLIT molecules bind to ROBO receptors through a highly conserved leucine-rich repeat (LRR) domain (52). In the AJBQM1 (*slit3^sa1569^)* line the loss of function mutation causes a truncation at amino acid 176 and in the *slit3^sa202^* line at amino acid 163. These are immediately adjacent to the LRR2 domain responsible for SLIT3’s functional interaction with ROBO proteins (52) and would therefore be predicted to lead to formation of non-functional proteins. Initially identified as a family of axon guidance molecules, SLIT proteins are known to be expressed in a range of tissues and, by regulating cell polarity, to play major roles in many developmental process including cell migration, proliferation, adhesion, neuronal topographic map formation and dendritic spine remodelling (53). *In vitro* SLIT proteins bind promiscuously to ROBO receptors suggesting that the proteins may co-operate *in vivo* in areas in which they overlap. However, their restricted spatial distributions, particularly of SLIT3 in the central nervous system (54) suggest the individual proteins play subtly different roles in vivo.

Despite its neuronal expression, the most prominent phenotype seen in SLIT3 deficient mice is postnatal diaphragmatic hernia (55, 56) with no obvious neuronal or axon pathfinding defects having been reported. Similarly, we did not detect any major differences in axon pathfinding nor in number of serotonergic and catecholaminergic cells in *slit3* mutant zebrafish larvae. As suggested previously (57) it may be that overlap of expression with other SLIT molecules compensates for loss of SLIT3 in the brain preventing gross neuronal pathfinding defects. However, subtle differences in circuit formation and/or axon branching may have escaped our analysis.

As our antibody staining may not have detected subtle, functionally important differences in dopaminergic and/or serotonergic circuit formation we examined the impact of the dopaminergic and serotonergic antagonist amisulpride on the acoustic startle response, a behaviour associated with vulnerability to addiction and known to be sensitive to modulation by dopaminergic antagonists in zebrafish as well as mammals (36,58,59). Although the binding affinity of amisulpride in zebrafish has not been examined, reported behavioural effects (60) are consistent with binding characteristics and behaviours seen in mammalian species. In mammals, amisulpride binds to D2/3 pre-and post-synaptic receptors with greater affinity at presynaptic than post-synaptic receptors (61, 62). Presynaptic D2 receptors in mammals act as autoreceptors and inhibit the synthesis and subsequent release of dopamine. D2/3 postsynaptic receptors are coupled to Gi/o G proteins mediating inhibitory neurotransmission (63). Treatment of rodents with D2/3 receptor antagonists leads to biphasic effects on locomotion such that low doses inhibit locomotion via pre-synaptic D2/3 autoreceptors, and high doses increase locomotion via post-synaptic D2/3 receptors (64). Treatment of adult zebrafish with amisulpride has a similar biphasic dose-dependent effect on locomotion such that low doses inhibit locomotion and high doses increase locomotion (60). These findings suggest that amisulpride has similar binding affinities at D2/3 receptors in fish as in rodents and imply the existence of pre-and post-synaptic D2/3 receptors with similar functional properties.

Our finding that high concentrations of amisulpride increased habituation to acoustic startle in wild type fish is in agreement with the effect of amisulpride in humans (36). A biphasic dose-dependent effect of amisulpride on habituation in wild type larvae suggests the involvement of both pre- and post-synaptic dopamine receptors. Inhibition of presynaptic receptors at low dose leading to increased responsiveness (reduced habituation), and inhibition of postsynaptic receptors at high doses causing reduced responsiveness (increased habituation). In contrast to results in wild type fish, *slit3^sa1569^* mutant larvae showed decreased habituation in the presence of both high and low dose amisulpride, suggesting a reduction of sensitivity to amisulpride at post-synaptic sites, possibly related to the marginal increase in dopamine D3 receptors in *slit3* mutants. Differential sensitivity to amisulpride was also seen at adult stages, where amisulpride inhibited nicotine-induced CPP in adult wild type fish but not in *slit3^sa1569^* mutants. These findings are consistent with a disrupted dopaminergic system, and/or disrupted interactions between dopaminergic and serotonergic systems caused by *slit3* loss of function.

Although our results are consistent with differences in dopaminergic signalling, differential sensitivity to actions of amisulpride at serotonergic receptors cannot be ruled out: amisulpride also acts as an antagonist at *Htr7* and *Htr2b* receptors with affinities approximately four times lower than at dopaminergic receptors. In mice, acoustic startle is sensitive to inhibition at *Htr2b* sites, and genetic ablation of the *Htr2b* receptors induced a reduction in startle amplitude and a deficit in prepulse inhibition of the startle reflex in loss of function mice (65). Loss of function of *Htr7* has no effect on acoustic startle or pre-pulse inhibition of acoustic startle in mice (66).

Although gene expression analyses revealed subtle up-regulation in several receptors - including *drd3, chrnb3, adra2b and htr4* - in *slit3* mutants, significant difference was only seen for the *ht1aa* receptor subtype. Zebrafish possess two homologues of the *HTR1A* gene, *htr1aa* and *htr1ab*, with overlapping expression domains (67). The observation of increased *htr1aa* expression in *slit3* mutants is of interest: Serotonergic signalling has been previously linked to drug reward processes including nicotine use and dependence (68, 69). Manipulations which decrease brain serotonin neurotransmission (e.g., a neurotoxic serotonin depletion or a lasting serotonin synthesis inhibition) elevate self-administration of several different drugs including nicotine in rats (69–71). Compounds that facilitate serotonin neurotransmission, such as selective serotonin reuptake inhibitors, decrease nicotine intake (72) whereas the HTR1A specific antagonist WAY100635 has also been reported to block nicotine enhancement of cocaine and methamphetamine self-administration in adolescent rats (73). Nicotine increases serotonin release in the striatum, hippocampus, cortex, dorsal raphe nuucleus (DRN), spinal cord and hypothalamus (74). The effects in the cortex, hippocampus, and DRN involve stimulation of *Htr1a* receptors, and in the striatum, *Htr3* receptors. In the DRN, *Htr1a* receptors play a role in mediating the anxiolytic effects of nicotine. In contrast, in the dorsal hippocampus and lateral septum, these same receptors mediate its anxiogenic effects. Further, pharmacological studies in rodents have shown that the *Htr1a* receptor antagonists WAY100635 and LY426965 alleviate the anxiety-related behavioural responses induced by nicotine withdrawal (75–77). Although it is possible that an anxiolytic effect of nicotine contributed to the increased nicotine-induced place preference, preliminary assessment of anxiety-like responses (tank diving) in *slit3* mutants, where mutants show decreased anxiety-like behaviour (n.s), argue against this.

It is perhaps of particular interest that it is a homologue of the HTR1A receptor that is up-regulated in *slit3* mutants. HTR1A is the major inhibitory serotonergic receptor in mammalian systems. In mammals, it is present as autoreceptors on cell bodies and dendrites in the raphe nucleus and as post-synaptic heteroreceptors in brain regions implicated in mood and anxiety such as prefrontal cortex, hippocampus and amygdala. Projections from raphe release serotonin throughout the entire forebrain. and brainstem and modulate a range of activities with additional raphe nuclei also providing innervation to the midbrain (see (78) for review). Interaction between HTR1A receptor and dopaminergic signalling are well established. Systemic administration of HTR1A receptor agonists leads to increased dopamine transmission in the nigrostrial pathway, ventral tegmental area and frontal cortex (78, 79). Although the mechanism underlying this increased dopamine transmission is not clear in all areas, the likely mechanism is by action on autoreceptors in the raphe so inhibiting serotonergic projections and disinhibiting dopaminergic transmission (79). Within the nucleus accumbens, a key area in drug reward, HTR1A agonists have little effect on dopamine release under baseline conditions, but inhibit amphetamine induced release (80). Further interactions between dopaminergic D2 receptor systems and HTR1A receptor signalling come from studies of atypical neuroleptics for the treatment of schizophrenia. Considerable evidence suggests that the balance between the properties of D2 receptors and HTR1A receptors influences the profile of action of these drugs in preclinical models (81, 82). Whilst synergistic effects of atypical neuroleptics may enhance dopamine release via inhibition of D2 autoreceptors and secondarily via disinhibition of projections from the raphe, it has been suggested (83) that association of D2 receptors and HTR1A in functional heterodimers that exhibit properties distinct to either G protein coupled receptor may also be involved. Such an interaction is seen between D2 receptors and adenosine A2a receptors both in vitro and in vivo and has been reported for D2 receptors and HTR1A in vitro (83). Whilst speculative, it is also potentially of relevance that HTR1A protein expression is upregulated in the brains of schizophrenics (84) and variants in both *HTR1A* and *SLIT3* are associated with psychiatric disorders (85–87), known to involve dopaminergic and serotoninergic pathways.

The mechanism by which loss of function in *slit3* leads to increased *htr1aa* expression and disrupted dopaminergic signalling in zebrafish mutants is yet to be established. However, *Htr1a* is up-regulated in conditions of reduced serotonergic signalling in other systems (88). Loss of serotonergic signalling from the DRN affects dopaminergic axonal outgrowth to the rat medial prefrontal cortex at developmental stages (89) such that lesioning of the DRN at neonatal stages results in significant increase in dopaminergic fibres. These effects are stage specific, raising the possibility that, although we did not detect any differences in catecholamine axon projections at three days post fertilisation, more detailed analysis at later stages of development would reveal significant differences. The observation that *slit3* is strongly expressed in the posterior raphe nucleus overlapping with *htr1aa* expression supports interaction between these two systems. Although detailed co-expression studies have not been performed, each of the other genes found to have marginal changes in expression also show overlapping expression with *slit3* (35,67,90,91).

Thus, our findings of an increase in *ht1aa* expression, altered sensitivity to amisulpride and altered nicotine CPP support a role for *slit3* signalling in the formation of dopaminergic and serotonergic pathways involved in responses to nicotine.

There are limitations to our findings: we used zebrafish of various ages in different experiments. While this confirms that loss-of-function in *slit3* alters behaviour from early life to adulthood, suggesting a developmental role, mechanisms underlying the two behavioural phenotypes may differ. We used whole embryos for the qPCR study so changes in expression in non-neuronal tissue may contribute to the observed differences, further, expression of genes in one tissues may mask changes of expression in another. In addition, we only examined a limited number of receptors and transporters for key neurotransmitter pathways. Important differences in other transmitter pathways and neurotransmitter metabolism may have been missed. We confirmed the translational effects of *SLIT3* gene variants in a human study, and the association was validated in a second, independent cohort. The sample sizes used in the human studies are small in comparison with those used in human discovery studies, however we used the two human studies to validate the findings in fish and analysed only a small set of SNPs focussed on just one gene which minimises the chance of type 1 error. Analyses were also corrected for multiple comparisons. The Finnish cohort was not used as a formal replication of the findings in the London cohort but instead to explore the effects of genetic variation on richer and more informative smoking phenotypes.

Associations between *SLIT3* and aspects of smoking phenotype have also been found in previous GWAS (https://atlas.ctglab.nl) (92). However, larger studies would be necessary to obtain greater precision on estimates of the effect size. Further studies are also required to determine the effects of genetic variation in *SLIT3* on anatomical pathways in the human brain and their functioning with view to identifying people who are at high risk of developing dependence. This could be achieved by using imaging techniques to study brain activation in response to smoking related cues in smokers who have the *SLIT3* polymorphisms linked to smoking (particularly rs12654448).

To our knowledge, this is the first report of a forward behavioural genetic screen in adult zebrafish successfully predicting a novel human coding genetic region involved in a complex human behavioural trait. Taken together, these results provide evidence for a role for *SLIT3* in regulating smoking behaviour in humans and confirm adult zebrafish as a translationally relevant animal model for exploration of addiction-related behaviours. Further work analysing the cellular processes affected as a result of the *Slit3* mutation may provide useful targets when designing tailored treatments to aid smoking cessation.

## MATERIALS AND METHODS

| Key Resources Table |  |  |  |  |
| --- | --- | --- | --- | --- |
| Reagent type (species) or resource | Designation | Source or reference | Identifiers | Additional information |
| Genetic reagent (*danio rerio*) | slit3 | Sanger Institute | sa202 | Generated by ENU mutagenesis. Soon to be available from ZIRC zebrafish resource centre |
| Genetic reagent (*danio rerio*) | slit3 | Sanger Institute | sa1569 | Generated by ENU mutagenesis. Soon to be available from ZIRC zebrafish resource centre |
| Strain (*danio rerio*) | Tupfel | Sanger Institute | Tubingen longfin | Wild type strain, now available from the ZIRC zebrafish resource centre |
| antibody | anti-5-HT (Rabbit polyclonal) | Sigma | Cat#S5545, | IHC (1/200) |
| antibody | anti-tyrosine hydroxylase (mouse monoclonal) | Abcam | Cat# AB152 | IHC (1/1000)) |
| Chemical compound | Amisulpride | Tocris | C2132 | 0.05-0.5mg/L |
| Chemical compound | varenicline | sigma | PZ0004 | 10-20micromolar |
| Chemical compound | bupropion | Sigma | B1277 | 1-10 micromolar |
| Chemical compound | Nicotine hemisulphate | Sigma | N1019 | 5-10 micromolar |

### Animals

All in vivo experimental work was carried out following consultation of the ARRIVE guidelines (NC3Rs, UK). Required sample size was estimated following pilot studies to determine effect sizes, and power calculations (beta = 0.8, alpha = 0.05). All animals were selected at random from groups of conspecifics for testing.

### Generation of F3 families of ENU-mutagenised fish

Wild type and ENU-mutagenized Tupfel longfin (TLF) fish were obtained from the Sanger Institute, as part of the Zebrafish Mutation Project which aimed to create a knockout allele in every zebrafish protein-coding gene [https://www.sanger.ac.uk/resources/zebrafish/zmp/]. At the Sanger, ENU-mutagenized TLF F_0_ males were outcrossed to create a population of F_1_ fish heterozygous for ENU-induced mutations. Due to the high ENU mutation rate (1/300 kb in F1 fish) and homologous recombination when F_1_ gametes are generated, all F_2_ were heterozygous for multiple mutations. F_2_ families, each generated from a separate F_1_ fish, were imported from the Sanger Institute to Queen Mary University of London (QMUL).

At QMUL a single male and female fish from each F_2_ family were inbred to generate 30 F_3_ families that would be 25% wild type, 50% heterozygous and 25% homozygous mutant for any single mutation, assuming Mendelian genetics. Based on exome sequencing data from the F_1_ generation performed at the Sanger, each F_3_ family contained 10-20 known predicted loss of function exonic mutations, approximately 100 non-synonymous coding mutations and approximately 1500 unknown mutations in non-coding domains across the entire genome (32). Breeding scheme is detailed in Figure 10.

**Figure 10.**
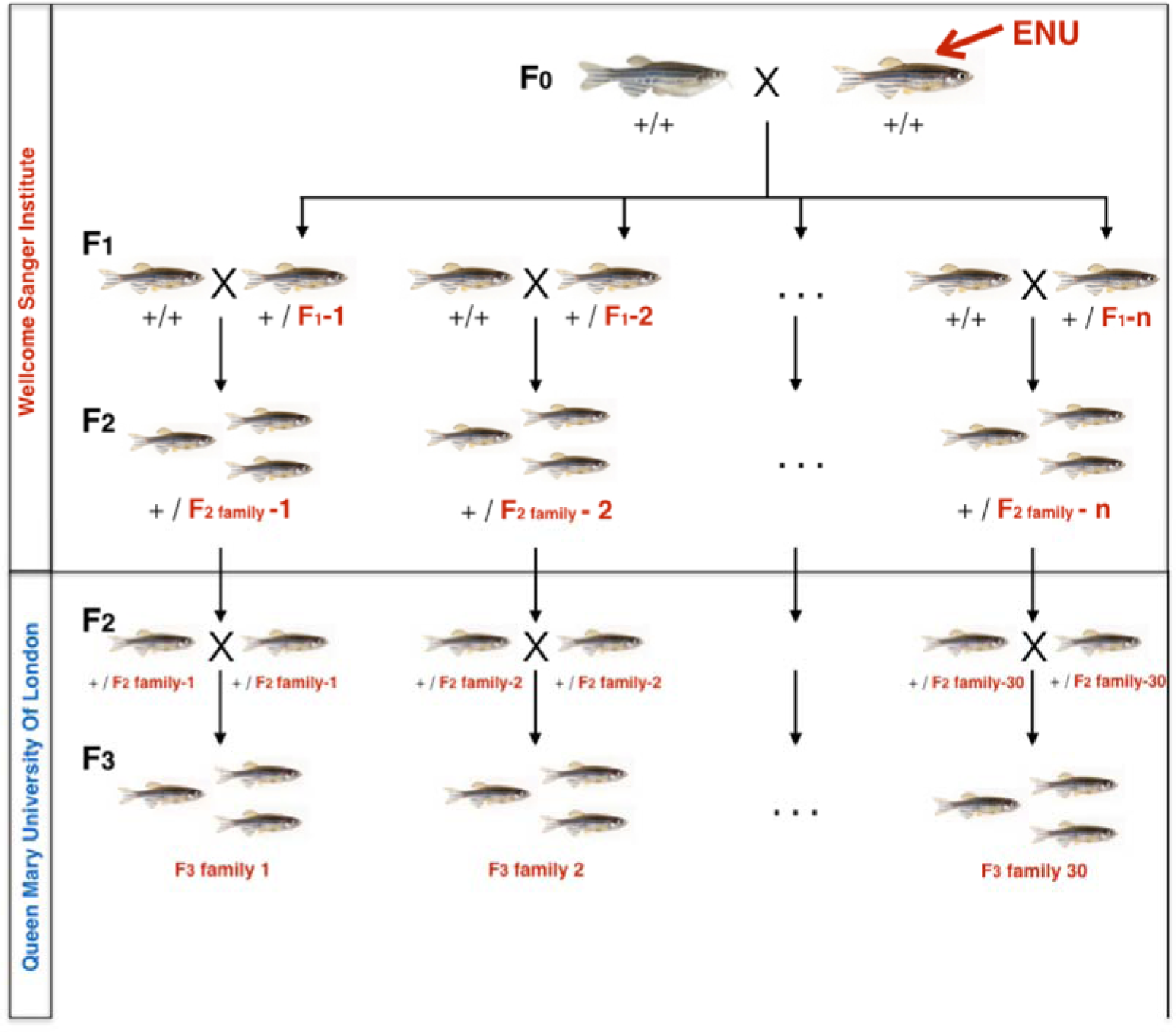
**Zebrafish breeding scheme to generate F3 families**. F_2_ ENU-mutagenized zebrafish, heterozygous for multiple mutations across the entire genome were obtained from the Wellcome Sanger Institute as part of the Zebrafish Mutation Project. At Queen Mary University of London, heterozygous F_2_ fish were incrossed to generate 30 F_3_ families, each containing 10-20 nonsense or essential splice site mutations and about 1500 additional exonic and intronic point mutations. F_3_ Families were arbitrarily numbered 1-30. Expected mutation rate and type of mutations in coding regions are specified on the right hand side.

### Fish maintenance

Fish were housed in a recirculating system (Tecniplast, UK) on a 14h:10h light:dark cycle (0830–2230). The housing and testing rooms were maintained at ∼25–28°C. Fish were maintained in aquarium-treated water and fed three times daily with live artemia (twice daily) and flake food (once). All procedures were carried out under license in accordance with the Animals (Scientific Procedures) Act, 1986 and under guidance from the local animal welfare and ethical review board at Queen Mary University of London.

### Conditioned place preference (CPP)

All fish were age and weight matched for all behavioural analysis and were approximately 5 months old, weighing 0.2-0.25g at the start of testing. Following habituation and determination of basal preference, animals were conditioned to 5μM nicotine (Sigma, Gillingham, UK Catalogue number: N1019) over three consecutive days and assessed for a change in place preference the following day. 5μM nicotine was used because it was predicted to induce a minimum detectable change in place preference based on results of previous studies (22, 24). This minimal effective dose was used to avoid possible ceiling effects if using a higher concentration. CPP was assessed as described previously (22,23,93): The testing apparatus was an opaque 3 L rectangular tank that could be divided in half with a Perspex divider. Each end of the tank had distinct visual cues (1.5cm diameter black spots versus vertical 0.5cm wide black and white stripes, matched for luminosity). After habituation to the apparatus and handling, we determined the basal preference for each fish: individual fish were placed in the tank for 10 min and the time spent at either end determined using a ceiling mounted camera and Ethovision tracking software (Noldus, Wageningen, NL). Any fish showing >70% preference for either end was excluded from further analysis (between 10 and 20% of fish). Fish were then conditioned with nicotine in the least preferred environment for 20 min, on 3 consecutive days: Each day each fish was restricted first to its preferred side for 20 min in fish water and then to its least preferred side with nicotine or, if a control fish, vehicle (fish water) added, for another 20 min. After 20 min in the nicotine (or vehicle)-paired environment each fish was returned to its home tank. After 3 days of conditioning, on the following day, fish were subject to a probe trial whereby each fish was placed in the conditioning tank in the absence of divider and the time spent at either end of the tank over a 10 min period was determined as for assessment of basal preference. The change in place preference was determined as the proportion time spent in the nicotine-paired zone during the probe trial minus the proportion time spent in the nicotine-paired zone during basal testing. The CPP procedure has been used and validated previously with nicotine as well as other drugs (22,23,93).

Data analysis: Change in preference scores were calculated as proportion time spent in drug paired stimulus after conditioning minus proportion time spent in drug paired stimulus before conditioning. Population means between generations were compared using independent two-sample t-tests, and effect-sizes ascertained using Cohen’s d (94). For the rescreen of outlier sibling families and *slit3^sa202^* line, mutant lines were compared with wild type controls using an independent two-sample t-test.

### CPP in the presence or absence of antagonists

To assess the ability of compounds (varenicline (Sigma, Gillingham, UK, PZ0004), buproprion (Sigma, Gillingham, UK, B1277) or amisulpride (Tocris, Bristol, UK, C2132) to inhibit subjective effects of nicotine a modified version of the CPP procedure was used (20). In this modified version, following habituation and establishment of basal preference, each day each fish was restricted first to its preferred side for 20 min in fish water and then removed from the conditioning tank and transferred to a tank containing the appropriate concentration of test compound or fish water (plus carrier where required) for 10min. After 10min the fish was returned to its least preferred side in the conditioning tank with nicotine or, if a control fish, vehicle (fish water) for another 20 min. After 20 min in the nicotine (or vehicle)-paired environment, each fish was returned to its home tank. After three days of conditioning to nicotine in the presence or absence of test compound, on the following day, fish were subject to a probe trial whereby each fish was placed in the conditioning tank in the absence of divider and the time spent at either end of the tank over a 10 min period determined. To assess the ability of varenicline (0-20μM) or buproprion (0-10μM) to inhibit subjective effects of nicotine in wild type fish, fish were incubated in the presence and absence of increasing doses of test compound (or vehicle) for 10 min before conditioning to 10μM nicotine. Statistical analysis was performed using a univariate analysis of variance (ANOVA), followed by Tukey’s post hoc test.

To test the effect of amisulpride on nicotine-induced CPP in wild type and *Slit3^sa1569^* mutant fish, fish were incubated in the presence or absence of 0.5mg/L amisulpride for 10 min before conditioning to 5μM nicotine. We selected 0.5mg/L amisulpride as this concentration has been shown to impact behaviour in adults previously (60) and wild type and homozygous mutants were found to be differentially sensitive to the effect of this concentration on acoustic startle response. Although previous studies with amisulpride in adult zebrafish used a pre-incubation period of 30min, to be consistent with our analysis of inhibitory effects of varenicline and bupropion, and to reduce the possibility of prolonged confinement in a small volume of drug affecting stress levels which may confound results, we used a preincubation period of 10 min.

Two-way ANOVA was performed with genotype (*slit3^+/+^ and slit3^sa1569/sa1569^*) and treatment (control, nicotine, nicotine+amisulpride) as independent variables. Values of p<0.05 were considered significant.

### Breeding and selection to assess heritability of nicotine-induced place preference

To test whether nicotine preference is heritable, fish falling in the upper and lower deciles of the ‘change in place preference’ distribution were kept for analysis and further breeding. Individuals were bred (in-cross of fish from the upper decile and in-cross of fish from the lower decile done separately) and their offspring screened for CPP (Second Generation CPP analysis). The same approach was repeated again: fish at the extremes of the Second Generation CPP distribution curve were selected and in-crossed, and their offspring were used to perform a Third Generation CPP analysis.

### Identification of ENU-induced mutations influencing nicotine place preference

To investigate whether ENU-induced mutations affect fish sensitivity to the rewarding effects of nicotine, candidate families were selected when the 3-4 fish from a family tested clustered together at one or other extreme of the change in preference distribution curve. To confirm the genetic effect on the CPP phenotype in candidate families, all remaining siblings of that family were assessed for nicotine induced CPP, along with non-mutagenized TLF control fish, to confirm the genetic effect on the phenotype. The experimental team and technical staff were blind to the fish genotype.

Candidate mutations, obtained from exome sequencing on F_1_ fish, were assessed for co-segregation with behaviour using site specific PCR (95) (See Supplementary Methods). Once a co-segregating candidate mutation was identified, larvae from an independent line carrying a predicted loss of function allele in the same gene were obtained from the Sanger Institute (*slit3^sa202^*) to confirm the association. Heterozygous *slit3^sa202/+^* and sibling *slit3^+/+^* larvae were reared to adulthood and assessed for nicotine-induced CPP as described above. All fish were fin clipped and genotyped following CPP.

### Characterization of larvae

#### Antibody staining

In order to visualize axonal pathways, fluorescent immunohistochemistry was carried out in three day old embryos from wild type *slit3^+/+^*, and homozygous mutant *slit3^sa1569/sa1569^* in-crosses. To prevent skin pigmentation, embryos were incubated in 0.2mM of 1-phenyl 2-thiourea (Sigma, Gillingham, UK) from 24 hours after fertilization. At three days, they were fixed in 4% paraformaldehyde (Sigma, Gillingham, UK) to avoid tissue degradation. For the immunostaining, rabbit polyclonal anti-tyrosine hydroxylase primary antibody (1:200; Sigma, Gillingham, AB152), rabbit polyclonal anti-serotonin (5HT) antibody (1:100, Sigma, Gillingham, S5545) and mouse anti-acetylated tubulin monoclonal antibody (1:1000; Sigma Gillingham, UK, T6793) were used. The three primary antibodies were detected with Alexa 546-conjugated secondary antibodies (1:400; Fisher Scientific, Loughborough, UK A11010). Whole-mount immunohistochemistry and mounting was performed as described previously (96).

#### Confocal microscopy imaging and analysis

Images were acquired using a Leica SP5 confocal microscope. Confocal z-stacks were recorded under the same conditions using diode laser and images were processed under ImageJ environment. Areas of interest for quantification were isolated, making sure that for all the individuals the same number of Z stacks, (covering the same D/V distance) were included. The number of cells was quantified using the ImageJ plugin “3D Objects counter” (https://biii.eu/3d-objects-counter). To quantify the number of catecholaminergic axons crossing the midline, a line was drawn from the Medulla oblongata interfascicular zone and vagal area to the locus coeruleus (Figure 5-figure supplement 1). Every 10 stacks (∼7.6 microns), the number of intensity peaks (defined as grey value intensity >20) was measured. Unpaired t-tests were calculated to assess genotype differences in the number of cells and intensity peaks.

#### Startle response in the presence or absence of amisulpride

Five day old larvae, generated from adult *slit3* wild type and homozygous mutant (*slit3^sa1569/sa1569^*) fish as for quantitative PCR, were individually placed in 24 well plates. In the drug-free condition, each well contained 300μl system water and 0.05% of dimethyl sulfoxide (DMSO, Sigma, Gillingham, UK). In the pharmacological conditions, serial dilutions of the dopaminergic and serotonergic antagonist amisulpride (Tocris, Bristol, UK, 71675-86-9) were prepared to give final concentrations of 0.05 mg/L, 0.1 mg/L or 0.5 mg/L amisulpride in 0.05% DMSO. Amisulpride concentrations were chosen based on previous studies in zebrafish (60) and correspond to 50, 100 and 500 times its Ki value for the D2 receptor in mammals (61, 62). To ensure that larvae were exposed to the drug for the same amount of time, amisulpride was added 15 minutes before undertaking the experiment. Care was taken regarding the distribution of concentrations and genotypes to ensure that experimental groups were randomly distributed in the plates. Plates were placed in a custom-made filming tower with a tapping device that applied 10 sound/vibration stimuli with two seconds interval between them. The setup for this device has been described elsewhere (97). Larval movement was recorded using Ethovision XT software (Noldus Information Technology, Wageningen, NL) and data were outputted in one second time-bins. Three technical replicates were performed (three different days) with three 24 well plates assayed each day.

For each fish, distance travelled (mm) during one second after each tap was recorded. All videos used to evaluate responses were checked for tracking errors. Any points where tracking errors were detected were removed. In total 105 points (31 wild type, 33 heterozygous, 41 homozygous) out of 5,460 were removed due to tracking errors. We also excluded individuals that failed to respond to all 10 taps using the criteria defined below. 39 wild type, 38 heterozygous, and 39 homozygous individuals were removed (a total of 115 individuals out of 546).

To evaluate how amisulpride dose and genotype affect habituation to the startle stimulus, we defined a threshold of distance moved: if a fish moved more than this threshold distance, we defined it as a ‘response’ (binary variable). We established our threshold by calculating the maximum distance each individual moved over all 10 taps. We ascertained that genotype was not a significant predictor of maximum distance through a linear model with maximum distance each individual moved as the response variable and genotype as the explanatory variable (ANOVA, F_2,543_ = 0.10, p=0.902). We therefore calculated the mean of the population maximum distance and defined our threshold as 70% of the mean maximum distance.

The percentage of fish responding to stimulus together with amisulpride dose, stimulus event number and genotype group were modelled in a beta regression conducted using the R package “betareg”. The proportion of individuals responding was the response variable and the three-way interaction between amisulpride dose, genotype, and stimulus event number was the explanatory variable. To determine whether this interaction was a significant predictor of individual responsiveness (indicating that genotypes varied in how amisulpride dose affected their habituation to repeated stimuli), likelihood ratio tests for nested regression models were performed. Results of all statistical analyses were reported with respect to a type-1 error rate of α=0.05.

#### Real-time quantitative PCR

Adult *slit3* wild type and *slit3^sa1569^* homozygous mutant fish, generated from a *slit3^sa1569/+^* heterozygous in cross, were bred to generate homozygous wild type, heterozygous mutant and homozygous mutant larvae. Embryos were carefully staged at 1, 24 and 48 hour and at five day post fertilisation to ensure, based on morphological criteria, there were no differences in development between groups. mRNA from 3 samples of five day old embryos (n=10 pooled embryos per sample) for each genotype was isolated using the phenol-chloroform method. cDNA was generated using the ProtoScript® II First Strand cDNA Synthesis Kit (NEB (UK Ltd.), Hitchen, UK). Relative qPCR assays were performed using the LightCycler 480 qPCR system from Roche Diagnostics, Ltd. with all reactions carried out in triplicates. Reference genes for all the qPCR analyses were β-actin, ef1α and rpl13α based on previous studies (93,98,99). Accession numbers and primer sequences for the genes can be found in Supplementary Table 2.

Relative mRNA expression in qPCR was calculated against reference gene cycle-threshold (Ct) values, and then subjected to one-way ANOVA. To account for multiple testing a Bonferroni correction was applied, and significance was declared at a threshold of 0.001.

#### Human Cohorts

In London human subjects were recruited from three clinical groups: patients with chronic obstructive pulmonary disease (COPD) (Cohort 1; n=272); patients with asthma (Cohort 2; n=293); and residents and carers in sheltered accommodation, with neither condition (Cohort 3; n=298). The methods used for recruitment and definition of phenotypes are reported elsewhere (100–102). The studies were approved by East London and The City Research Ethics Committee 1 (09/H0703/67, 09/H0703/76 and 09/H0703/112). Written informed consent was obtained from all participants.

Details of the Finnish twin cohort are reported elsewhere (103–105). In brief, twin pairs concordant for moderate to heavy smoking were identified from the population-based Finnish Twin Cohort survey responders. The twin pairs and their siblings were invited to a computer-assisted, telephone-based, structured, psychiatric interview (SSAGA) (103), to yield detailed information on smoking behaviour and nicotine dependence as defined by Fagerström Test for Nicotine Dependence (FTND) and DSM-IV diagnoses. Human phenotypes to be investigated in relation to zebrafish nicotine seeking behaviour were determined by consensus *a priori*.

Sample characteristics of the human cohorts and detailed definitions of both London and Finnish phenotypes can be found in the supplementary material and Supplementary Table 3.

#### Human genotyping

For the London cohorts, DNA from participants was extracted from whole blood using the salting-out method (106) and normalized to 5ng/µl. 10ng DNA was used as template for 2 µl TaqMan assays (Applied Biosystems, Foster City, CA, USA) performed on the ABI 7900HT platform in 384-well format and analysed with Autocaller software. Pre-developed assays were used to type all SNPs. See

Supplementary Table 4 for primer and reporter sequences. Typing for two SNP (rs6127118 and rs11574010) failed. For the Finnish cohort, DNA was extracted from whole blood and genotyping was performed at the Wellcome Trust Sanger Institute (Hinxton, UK) on the Human670-QuadCustom Illumina BeadChip (Illumina, Inc., San Diego, CA, USA), as previously described (103–105). Genotyping and imputation for the Finnish cohort at the Wellcome Trust Sanger Centre have been described previously (107).

#### Human association analyses

We attempted to replicate the zebrafish findings initially in a cohort from London using a narrow set of SNPs in *SLIT3*. We then used the same set of SNPS to evaluate effects on more detailed smoking phenotypes in a Finnish twin cohort.

London cohort association analysis was performed using PLINK v1.07 (108). *SLIT3* SNPs that had been previously associated with disease phenotype were identified and the 20 with low linkage disequilibrium score selected for analysis. Of twenty *SLIT3* SNPs, one departed from Hardy-Weinberg equilibrium (rs13183458) and was excluded. Linear regression was performed on average number of cigarettes smoked per day, controlling for age, sex and cohort. Since physiological and genetic mechanisms may be different in heavy (more dependent) and light (less dependent) smokers we repeated on heavy smokers (≥ 20 cigarettes per day) and light smokers (< 20 cigarettes per day). Smoking cessation (current *vs* ever smokers) was analysed using logistic regression controlling for age, sex and cohort. All analyses were performed under the additive genetic model and multiple testing was taken into account using the Benjamini-Hochberg adjustment. Only individuals from European ancestry were included in analyses.

Association analyses for the Finnish Twin Cohort were performed using GEMMA v0.94 (109) with linear mixed model against the allelic dosages controlling for age and sex. Sample relatedness and population stratification were taken into account by using genetic relatedness matrix as random effect of the model.

## Acknowledgements

Funding: MRC UK, G1000403 (CHB/RW). NC3Rs G1000053 (CHB). BBSRC BB/M007863 (CHB) NIH U01 DA044400-03 (CHB) NIHR PGfAR RP-PG-0609-10181 (RW). NIHR PGfAR, RP-PG-0407-10398 (ARM). CHB is a member of the Royal Society Industry Fellows’ College. RW is an NIHR Senior investigator (NF-SI-0515-10076). VK holds a Wellcome Trust Clinical Research Fellowship. JK is supported by the Academy of Finland (grants 308248, 312073). CHB conceived the project and directed the research. RW, AM and JK oversaw the human studies. AJB, JGG, DJ, MTT, AS, MOP, CHB conducted experiments. TP, VK analysed the human data. EMB and DS supplied the mutant lines. Some of the data in this manuscript has been published previously as a preprint: bioRxiv 453928: doi: https://doi.org/10.1101/453928

## Supplementary Methods

### Zebrafish genomic DNA extraction

Genomic DNA was extracted from fin-clips using QIAGEN DNeasy® Blood and Tissue Kit (Qiagen, Manchester, UK) according to manufacturer’s instructions. Samples were eluted into distilled water and stored at −20°C until later use.

### Site Specific Polymerase Chain Reaction

Allele-specific PCR single nucleotide polymorphism (SNP) assays were used for genotyping F3 individuals for mutations known to be present in the ENU-mutagenized F1 generation. Four primer pairs were designed to carry out PCR genotyping as previously described (95). The list of loss-of-function mutations in the AJBQM1 and AJBQM2 lines is detailed in Supplementary Table 1. For each line, a primer was designed with 3’ complementary to the ENU-SNP with a second primer ∼100bp downstream. The second pair had one primer with 3’ complementary to the wild type base with a second primer ∼200bp upstream. The resulting PCR results in a 300bp fragment that spans the region and acts as an internal control for the PCR plus one 100bp fragment if homozygous for the mutation, 2 bands of 100bp and 200bp if heterozygous, and one 200bp fragment if homozygous wild type. The 4-primer groups were designed with melting temperatures as close as possible using the NCBI primer design tool and were ordered from Eurofins, MWG operon (Ebersberg, DE).

### Human Association Studies

#### Phenotype definitions for the London cohorts

*<u>Amount smoked</u>* was defined as the average number of cigarettes smoked per day (CPD) for each participant. Participants met criteria for *<u>smoking cessation</u>* if they reported being ‘ever smokers’ and reported ***not*** smoking currently. The percentage of current smokers in the cohort was 42%, 7% and 18% for ViDiCO, ViDiAs and ViDiFLU, respectively.

#### Phenotype definition for the Finnish twin cohort study

Definitions of the phenotypes were adapted from Broms et al (105).

#### Amount smoked

Cigarettes per day (CPD) constitutes of eight categories: 1-2, 3-5, 6-10, 11-15, 16-19, 20-25, 26-39, ≥40 CPD. In the statistical analyses of the CPD variables, original categorical observations were replaced with class means of CPD (1.5, 3.5, 8, 13, 17.5, 22.5, 32.5, and 45 cigarettes per day, respectively). Regression coefficients can therefore be interpreted as the average change in number of cigarettes smoked per day when the number of minor allele is increased by one.

⍰***CPD***: Number of cigarettes smoked per day during month of heaviest smoking. Values ranged from 1 to >40 with mean=19.8 cigarettes per day.
⍰***Maximum CPD***: Maximum number of cigarettes ever smoked during one day (24h period). Values ranged from 2 to 98 with mean=30 cigarettes per day.

#### Smoking initiation

⍰***Age of onset of weekly smoking:*** Age (years) when started to smoke weekly (“How old were you when you first smoked a cigarette at least once a week for at least two months in a row?”). Values ranged from 6 to 54, mean=17.3 years.
⍰***First time sensation.*** Sensation felt after smoking the first cigarette or first puffs. Sensation measured as: “While smoking your very first cigarettes, did you (1) like the taste or smell of the cigarette, (2) cough, (3) feel dizzy or light-headed, (4) feel more relaxed, (5) get a headache, (6) feel a pleasurable rush or buzz, (7) feel your heart racing, (8) feel nauseated, like vomiting, (9) feel your muscles tremble or become jittery, (10) feel burning in your throat”). Sum score of 10 questions (items #1, #4, and #6 were reverse-scored before summation): 0 points if answered “No”, 1 = “A little bit”, 2=”Some”, 3= “Quite a bit”, 4=”A great deal”. Cronbach’s alpha = 0.70. Values ranged from 3.6 to 15.8. Mean =10.2.

#### Nicotine dependence

⍰***DSM-IV ND diagnosis***: Nicotine dependence by DSM-IV diagnosis (≥3 symptoms out of 7 occurring within a year). Prevalence = 53.5%.
⍰***DSM-IV ND symptoms***: Number of DSM-IV ND symptoms from 0 to 7. Mean=3
⍰***FTND (***≥***4):*** Nicotine dependent if ≥4 out of 10 points in Fagerström Test for Nicotine Dependence. Prevalence = 50.4%
⍰***FTND score:*** Fagerström Test for Nicotine Dependence (FTND) score: 0 to 10 points. Mean=3.7.
⍰***FTND time to first cigarette (TTF):*** Time to first cigarette in the morning (one item of the FTND scale). Five categories: 0-5 min, 6-15 min 6, 16-30 min 6, 31-60 min, >60 min. Categorization differs from original four categories (3), i.e., 6-30 minutes is split into 6-15 min and 16-30 min. In our data set 46% of smokers belong to the group of 6-30 min, and from the smoking behaviour point of view there is a significant difference whether one smokes the first cigarette within 6 minutes or 30 minutes from waking up. In this data set 22% of smokers belong to the 6-15 min and 24% to the 16-30 min group. Values ranged from 1 to 5 with a mean=3.1.

## Supplementary results

**Figure 5-figure supplement 1.**
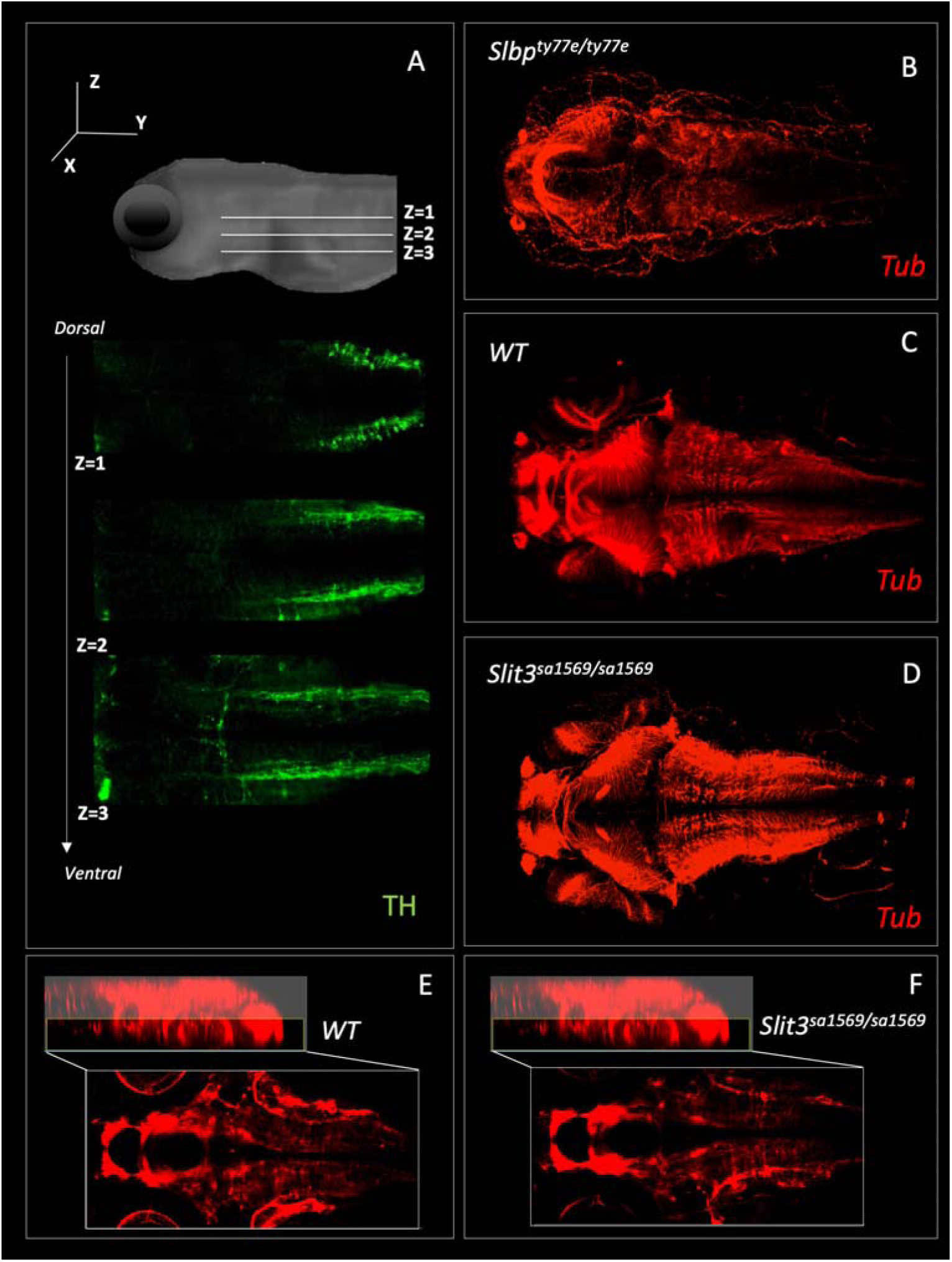
Fluorescent immunohistochemistry in three-day old wild type and homozygous mutant *slit3^sa1569^* labelled with tyrosine hydroxylase (A) and tubulin (B-F). A) Example of three Z-planes used to quantify catecholaminergic projections crossing the midline. (Stacks 1-3). B) Anti-tubulin staining of *slbp^ty77e/ty77e^* mutant larvae, known to have fewer neurons and axonal defects (110). C-D) Anti-tubulin staining of wild type and homozygous mutant *slit3^sa1569^*. E-F) Zoom-in of C-D for midline in the ventral forebrain, where *slit3* is known to be expressed (35).

**Figure 6 –figure supplement 1.**
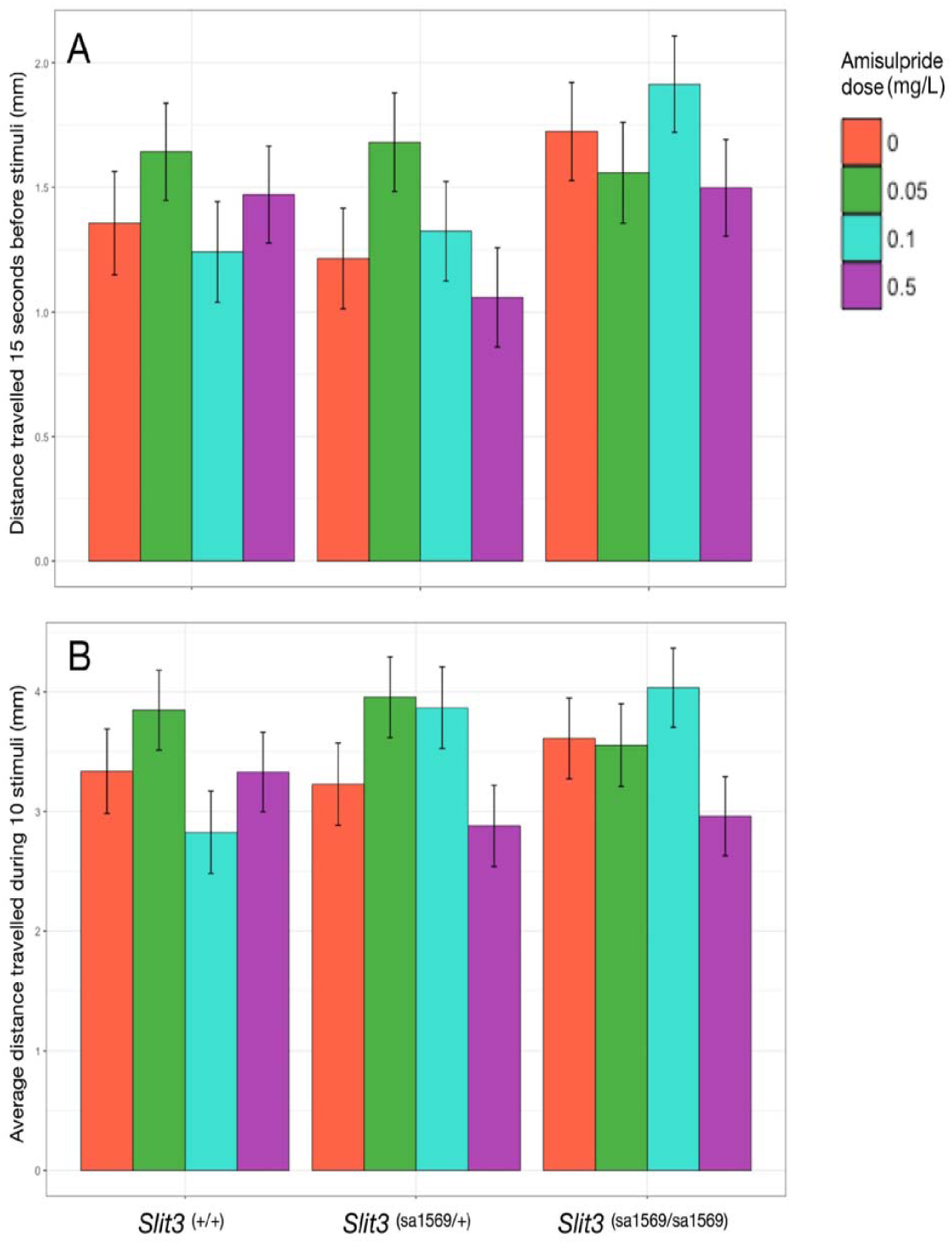
Average distance moved before (Figure 1A) and during startle stimuli (Figure 1B) in wild type and *slit3^sa1569^* mutant five day old zebrafish larvae. **A)** Distance moved as function of amisulpride dose, fish *slit3* genotype and their interaction. The effect of dose and genotype was tested in a linear mixed model. Timepoint, well where the fish were placed and plate were also included as fixed factors and the fish ID as random factor. **B)** Distance moved during taps as function of amisulpride dose and fish *slit3* genotype. Drug and genotype effects were examined in a linear mixed model including stimulus number, well, plate used and distance moved before stimuli as fixed factors and Fish ID as random factor. Zebrafish larvae did not differ in the distance travelled before or during startle stimuli as a function of amisulpride dose nor genotype (p > 0.05). Bars represent estimated marginal means ± SEM (n=42-48 fish per group).

**Supplementary Table 1:**
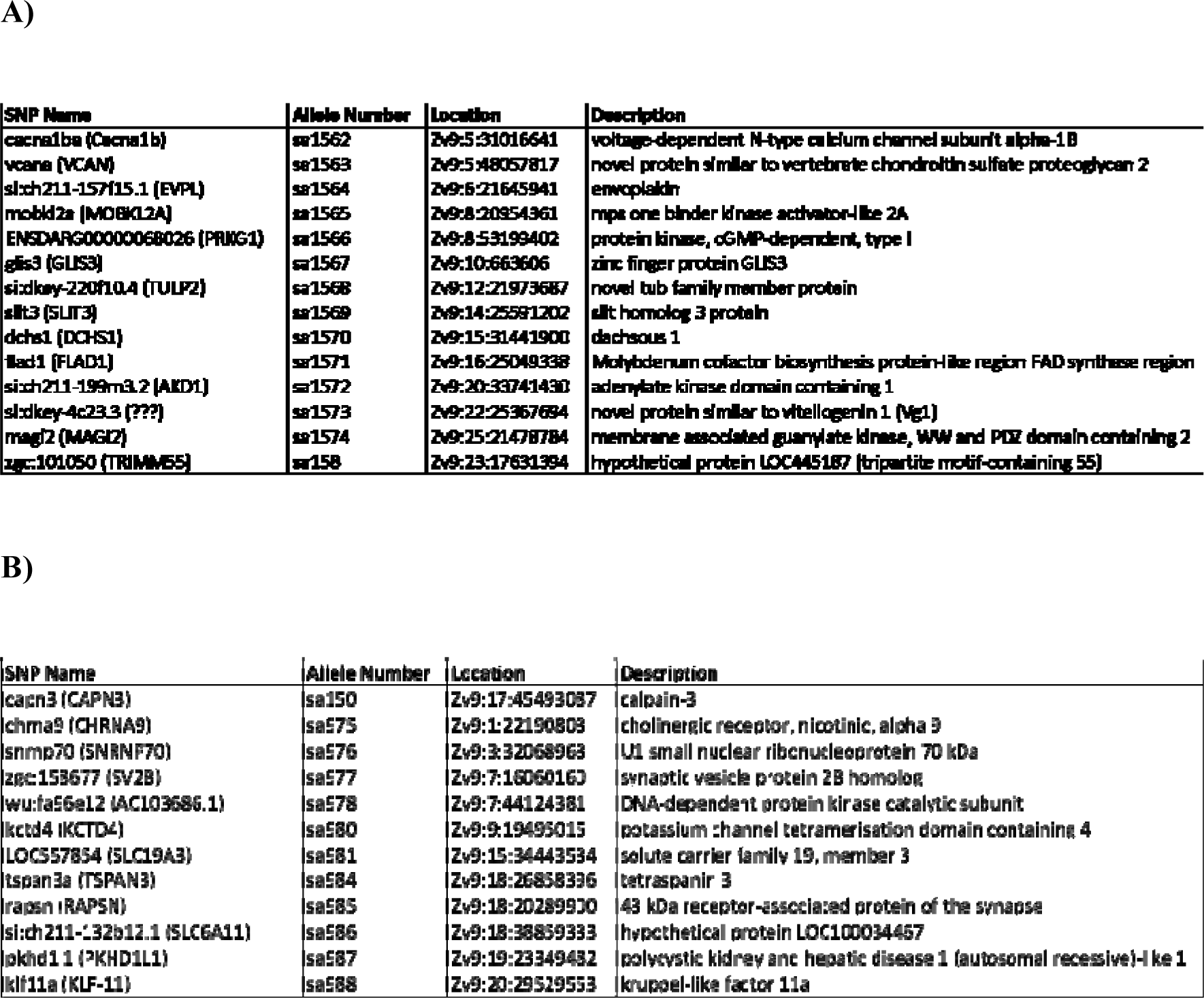
List of loss-of-function mutations in the AJBQM1 (A) and AJBQM2 (B) lines. List was derived from exome sequencing and provided by the Wellcome Sanger Trust, Hinxton, Cambridge.

**Supplementary Table 2.**
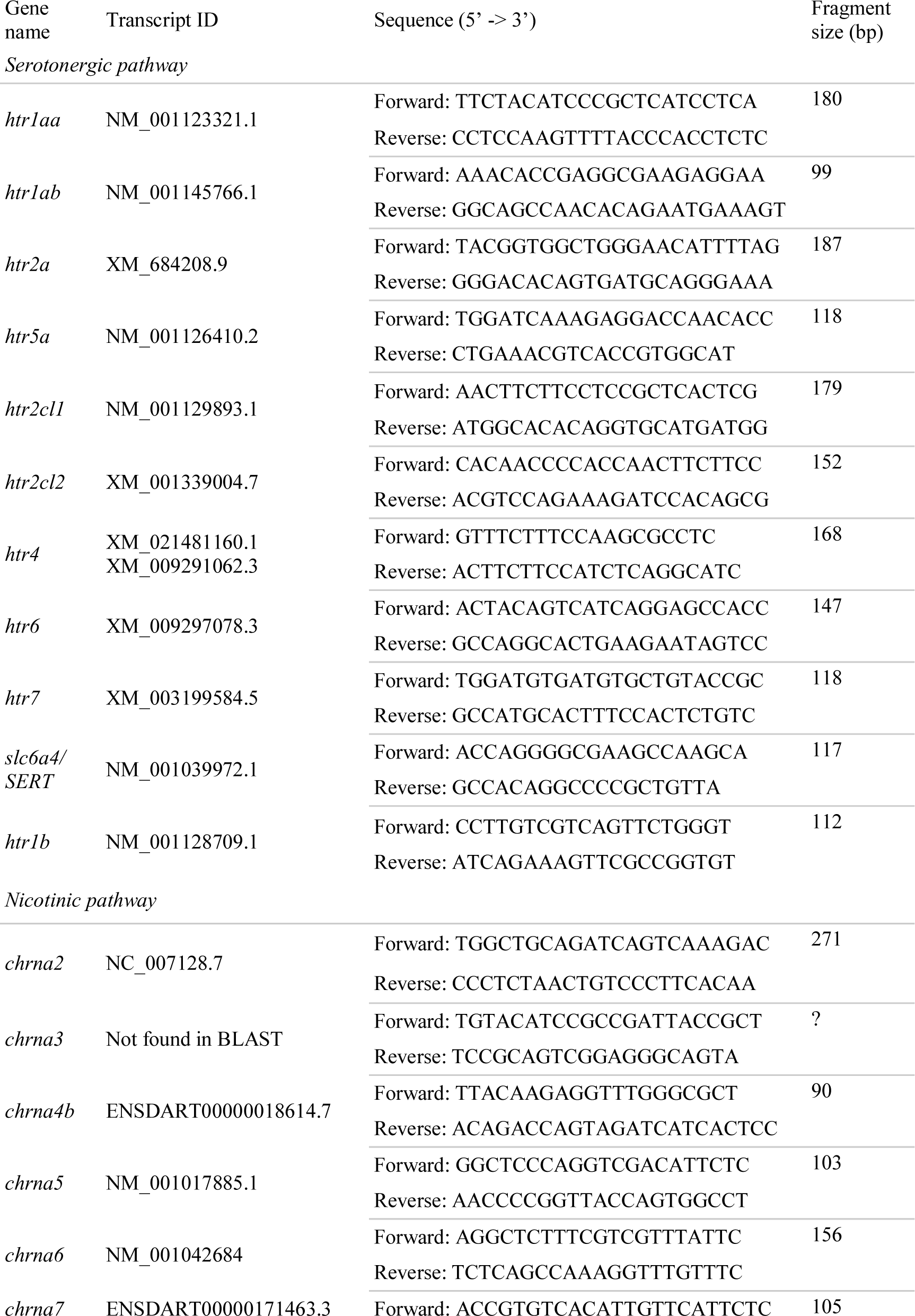
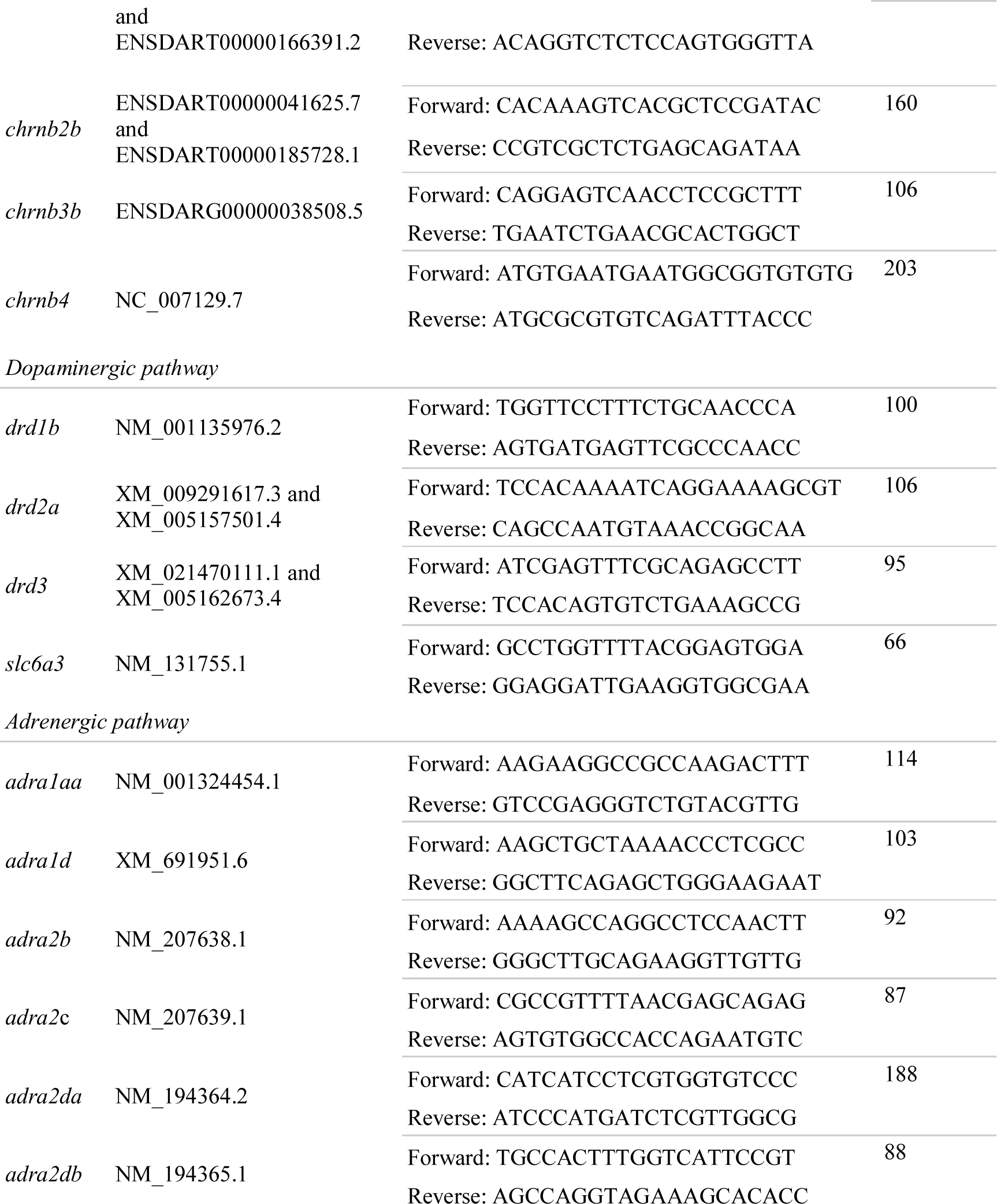
Gene ID and primer sequences used in gene expression analysis.

**Supplementary Table 3:**
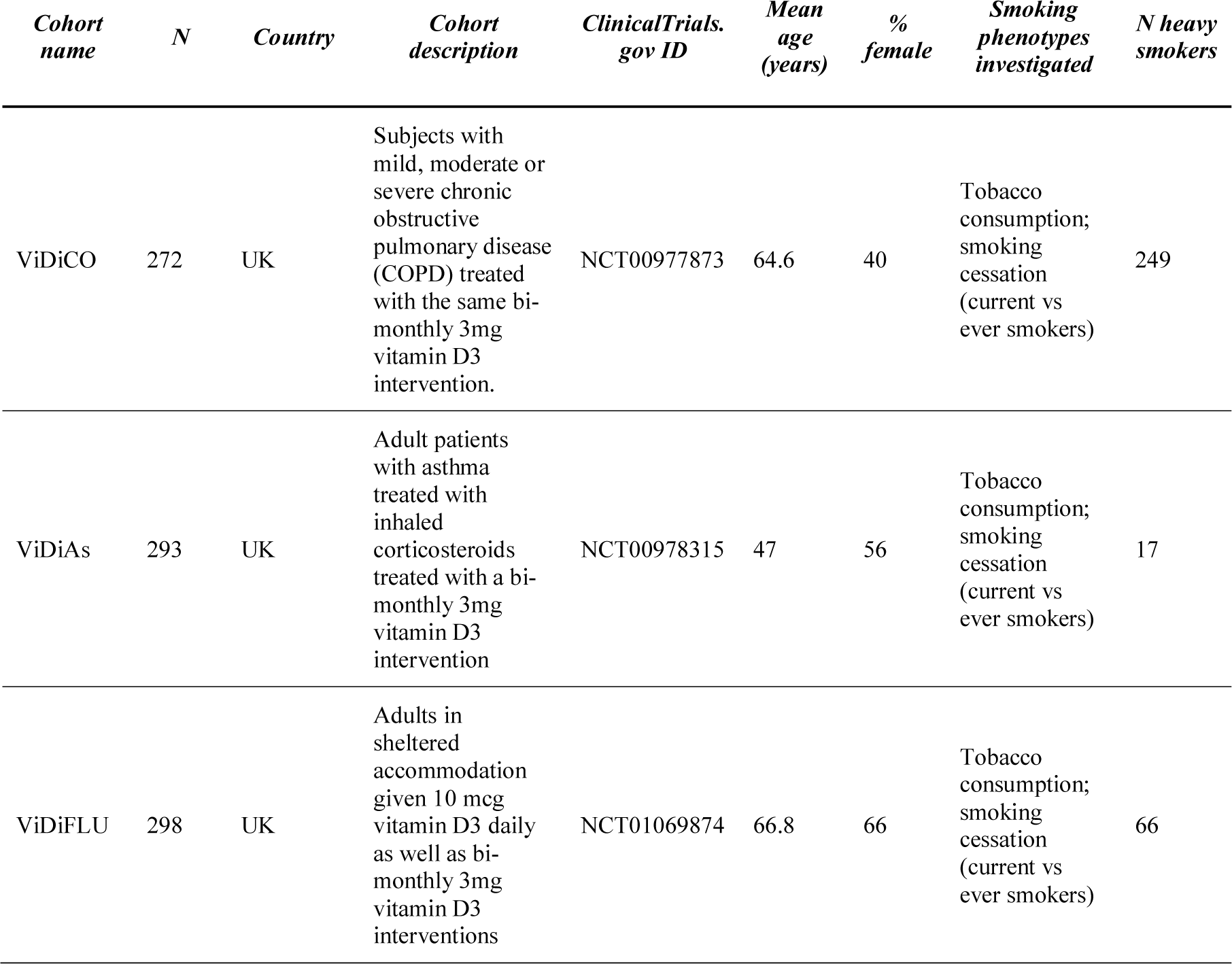
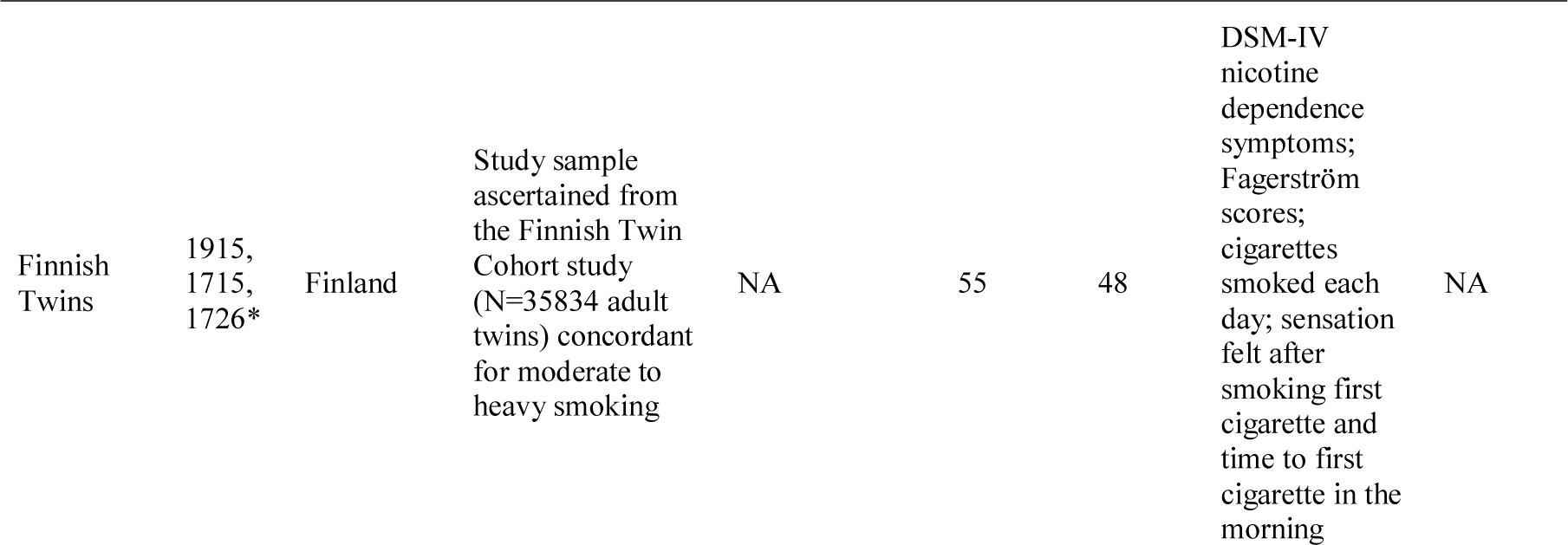
**Sample characteristics of the human cohorts.** Detailed inclusion and exclusion criteria for the London cohorts can be found in https://clinicaltrials.gov. Further details about recruitment, power calculation and definition of medical phenotypes can be found elsewhere (100–102). Genotype and phenotype Finnish twins are deposited in the Biobank of the National Institute for Health and Welfare (https://thl.fi/en/web/thl-biobank/for-researchers). **Differences in sample size for Finnish cohorts was due to hard-call genotype probability threshold. DSM-IV nicotine dependence symptoms, Fagerström scores and cigarettes smoked each day (N = 1715). Sensation felt after smoking first cigarette (N = 1915). Time to first cigarette in the morning (N= 1726).

| *Cohort name* | *N* | *Country* | *Cohort description* | *ClinicalTrials.gov ID* | *Mean age (years)* | *% female* | *Smoking phenotypes investigated* | *N heavy smokers* |
| --- | --- | --- | --- | --- | --- | --- | --- | --- |
| ViDiCO | 272 | UK | Subjects with mild, moderate or severe chronic obstructive pulmonary disease (COPD) treated with the same bi-monthly 3mg vitamin D3 intervention. | NCT00977873 | 64.6 | 40 | Tobacco consumption; smoking cessation (current vs ever smokers) | 249 |
| ViDiAs | 293 | UK | Adult patients with asthma treated with inhaled corticosteroids treated with a bi-monthly 3mg vitamin D3 intervention | NCT00978315 | 47 | 56 | Tobacco consumption; smoking cessation (current vs ever smokers) | 17 |
| ViDiFLU | 298 | UK | Adults in sheltered accommodation given 10 mcg vitamin D3 daily as well as bi-monthly 3mg vitamin D3 interventions | NCT01069874 | 66.8 | 66 | Tobacco consumption; smoking cessation (current vs ever smokers) | 66 |
| Finnish<br>Twins | 1915,<br>1715,<br>1726* | Finland | Study sample<br>ascertained from<br>the Finnish Twin<br>Cohort study<br>(N=35834 adult<br>twins) concordant<br>for moderate to<br>heavy smoking | NA | 55 | 48 | DSM-IV<br>nicotine<br>dependence<br>symptoms;<br>Fagerström<br>scores;<br>cigarettes<br>smoked each<br>day; sensation<br>felt after<br>smoking first<br>cigarette and<br>time to first<br>cigarette in the<br>morning | NA |

**Supplementary Table 4.**
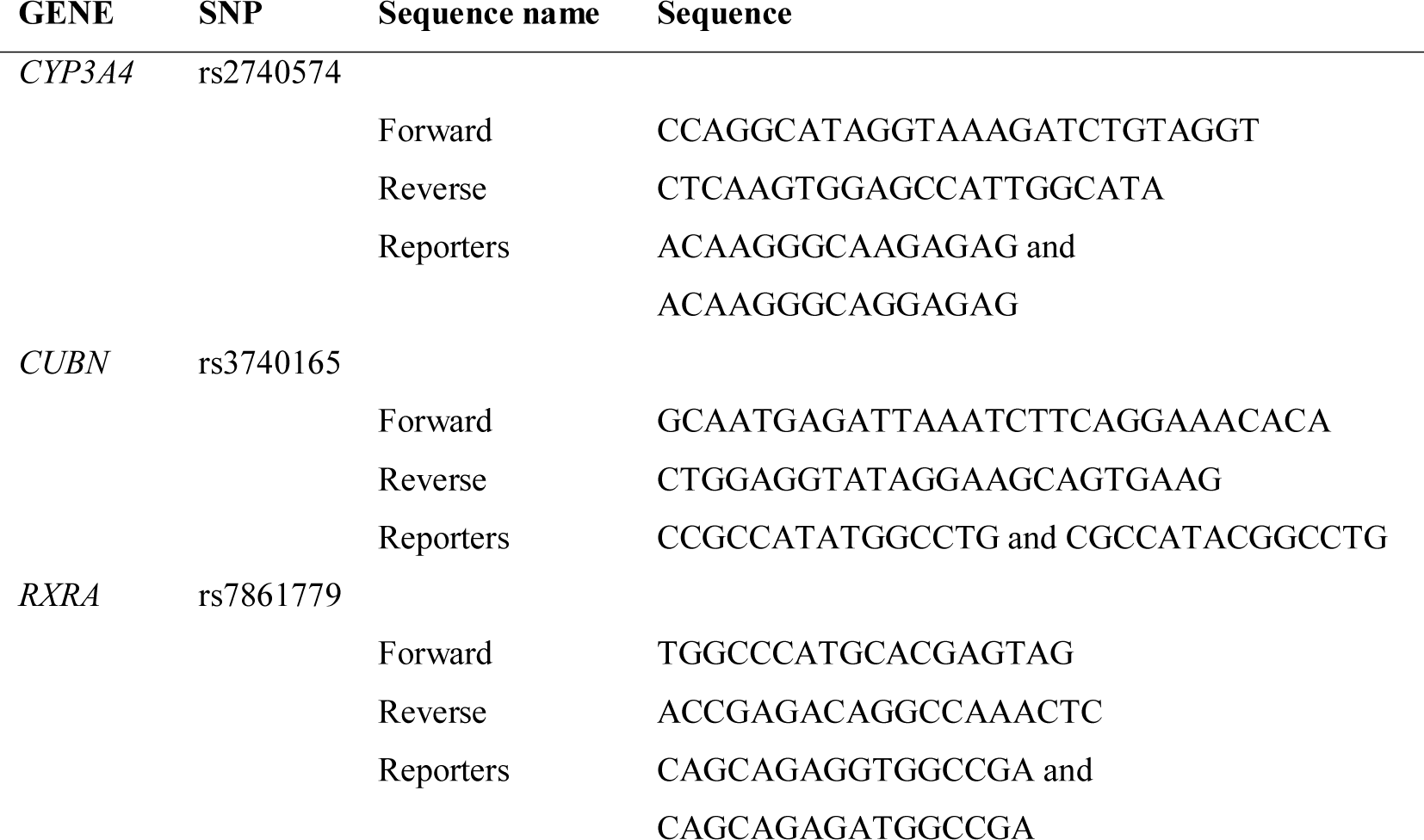
Primer and reporter sequences used for human genotyping.

**Supplementary Table 5:**
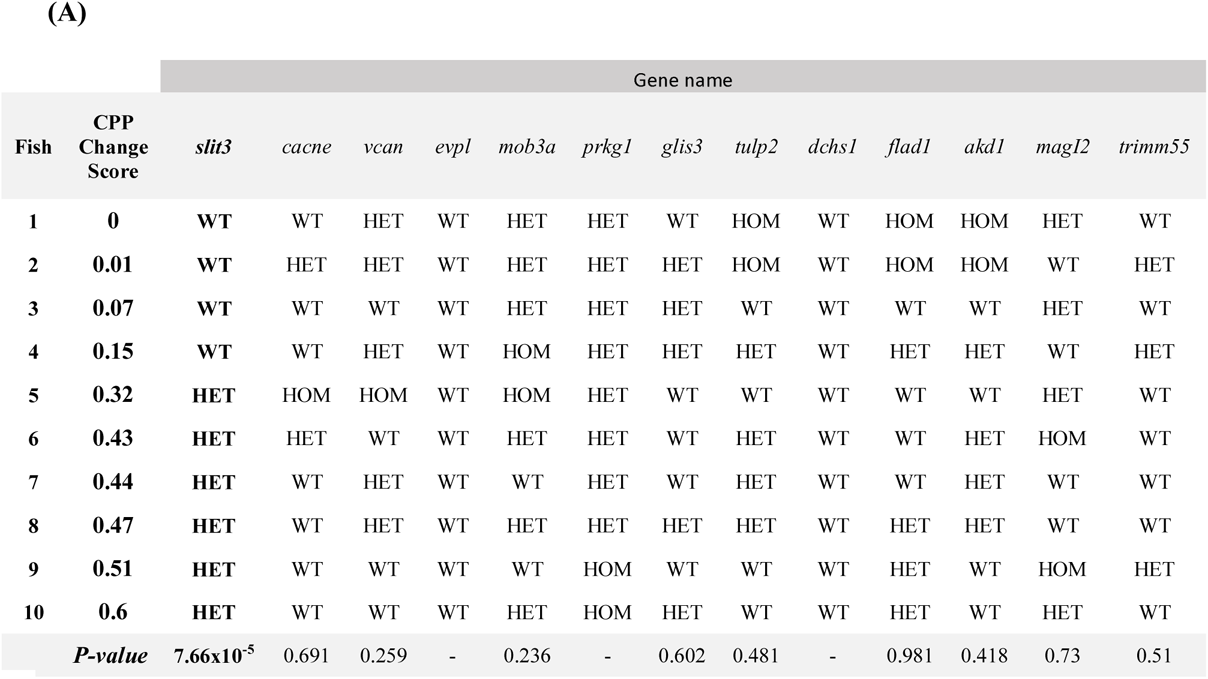
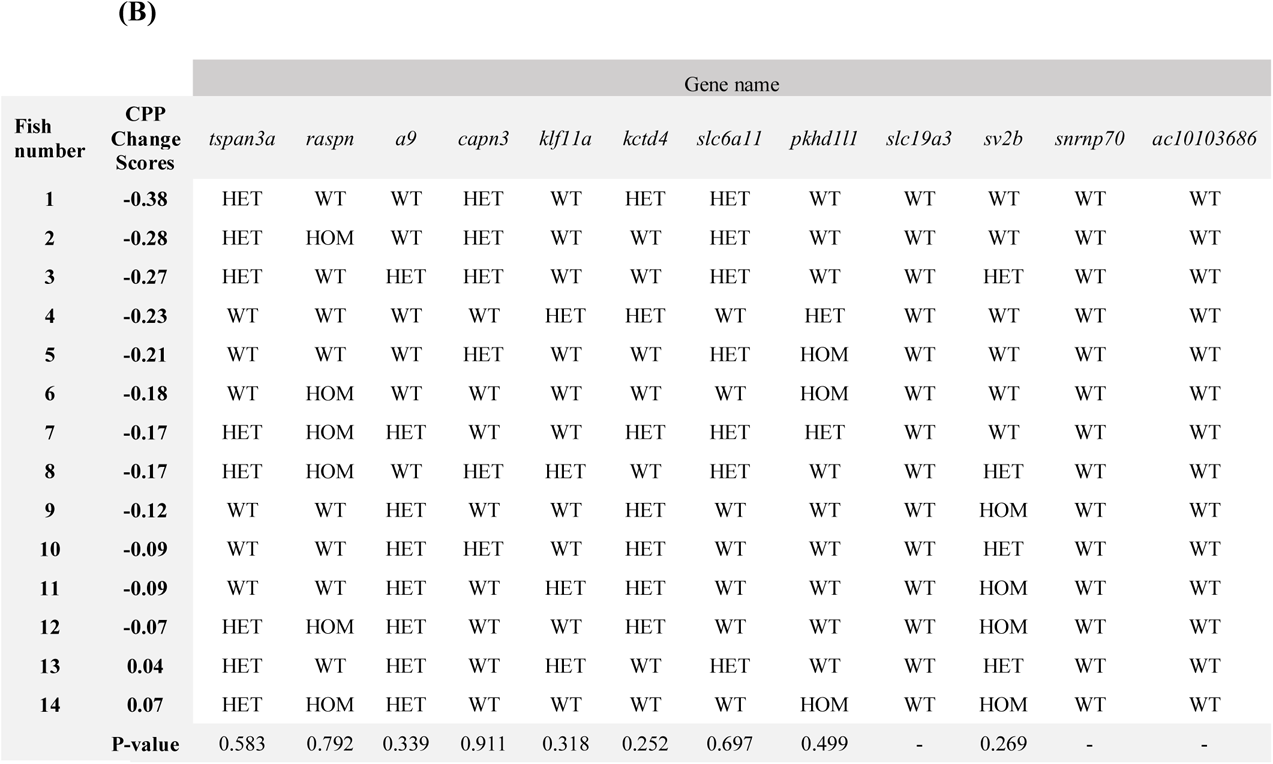
Site specific PCR genotyping of AJBQM1 (A) and AJBQM2 **(B) outlier siblings.** The siblings were genotyped at each of the candidate loci using site specific PCR and results compared with each individual place preference change scores. Each row represents one fish, with their corresponding CPP change score, and genotype for each locus assessed (13 loci for AJBQM1, 12 loci for AJBQM2). P-values result from independent two-sample t-tests comparing the CPP change scores between genotype groups at each locus. **(A)**

**Supplementary Table 6:** Results of association analysis of *SLIT3* SNPs on **smoking initiation.** Logistic regression of initiation vs non-initiation on additive genotype, controlling for age, sex and cohort. OR: Odds ratio. >1 value indicates that the minor allele increases odds of persistent smoking relative to the major allele, SE: standard error, L95: lower limit of 95% confidence interval, U95: upper limit of 95% confidence interval. Benjamini Hochberg cut off at 0.1 = 0.00526.

| **SNP** | **OR** | **SE** | **L95** | **U95** | **P value** |
| --- | --- | --- | --- | --- | --- |
| **rs2938774** | **0.7253** | **0.1418** | **0.5493** | **0.9578** | **0.02357** |
| **rs11742567** | **1.328** | **0.1538** | **0.9825** | **1.796** | **0.06496** |
| **rs4282339** | **0.7277** | **0.1815** | **0.5099** | **1.039** | **0.07991** |
| **rs297886** | **1.328** | **0.176** | **0.9405** | **1.875** | **0.1071** |
| **rs9688032** | **1.269** | **0.1495** | **0.9467** | **1.701** | **0.111** |
| **rs7728604** | **1.198** | **0.1446** | **0.9024** | **1.591** | **0.2112** |
| **rs1345588** | **0.7788** | **0.2046** | **0.5215** | **1.163** | **0.2218** |
| **rs17734503** | **0.7362** | **0.2632** | **0.4395** | **1.233** | **0.2445** |
| **rs12515725** | **0.8612** | **0.1458** | **0.6471** | **1.146** | **0.3052** |
| **rs11749001** | **0.8698** | **0.1951** | **0.5933** | **1.275** | **0.4746** |
| **rs3733975** | **1.116** | **0.1641** | **0.8092** | **1.54** | **0.5029** |
| **rs12521041** | **1.116** | **0.1641** | **0.8092** | **1.54** | **0.5029** |
| **rs11134527** | **0.9212** | **0.155** | **0.6798** | **1.248** | **0.5964** |
| **rs12654448** | **0.8718** | **0.2654** | **0.5182** | **1.467** | **0.6052** |
| **rs1559051** | **0.9257** | **0.1575** | **0.6799** | **1.26** | **0.6241** |
| **rs17665158** | **0.9304** | **0.171** | **0.6654** | **1.301** | **0.673** |
| **rs1421763** | **0.9423** | **0.1704** | **0.6748** | **1.316** | **0.7271** |
| **rs295994** | **0.9732** | **0.1404** | **0.7391** | **1.281** | **0.8464** |
| **rs10036727** | **1.01** | **0.1522** | **0.7491** | **1.361** | **0.9503** |

**Supplementary Table 7:** Results of association analysis of *SLIT3* SNPs on **persistent smoking.** Logistic regression of initiation vs non-initiation on additive genotype, controlling for age, sex and cohort. OR: Odds ratio. >1 value indicates that the minor allele increases odds of persistent smoking relative to the major allele, SE: standard error, L95: lower limit of 95% confidence interval, U95: upper limit of 95% confidence interval. Benjamini Hochberg cut off at 0.1 = 0.00526.

| **SNP** | **OR** | **SE** | **L95** | **U95** | **P value** |
| --- | --- | --- | --- | --- | --- |
| rs11134527 | 1.428 | 0.1573 | 1.049 | 1.943 | 0.02359 |
| rs12521041 | 0.6871 | 0.1736 | 0.489 | 0.9655 | 0.03061 |
| rs11742567 | 0.7288 | 0.1547 | 0.5382 | 0.987 | 0.04089 |
| rs3733975 | 0.7165 | 0.1712 | 0.5123 | 1.002 | 0.05146 |
| rs17734503 | 0.6142 | 0.2631 | 0.3667 | 1.029 | 0.06398 |
| rs1345588 | 0.6786 | 0.2145 | 0.4457 | 1.033 | 0.07068 |
| rs17665158 | 1.338 | 0.163 | 0.9722 | 1.842 | 0.07393 |
| rs12654448 | 0.6225 | 0.2671 | 0.3688 | 1.051 | 0.07597 |
| rs2938774 | 1.232 | 0.1394 | 0.9373 | 1.619 | 0.1348 |
| rs295994 | 1.214 | 0.1443 | 0.9147 | 1.61 | 0.1796 |
| rs7728604 | 1.152 | 0.1434 | 0.8699 | 1.526 | 0.3232 |
| rs1559051 | 1.115 | 0.1549 | 0.8231 | 1.511 | 0.4821 |
| rs12515725 | 0.9108 | 0.1456 | 0.6846 | 1.212 | 0.521 |
| rs297886 | 0.9342 | 0.1726 | 0.6661 | 1.31 | 0.6932 |
| rs4282339 | 0.9391 | 0.19 | 0.6471 | 1.363 | 0.7411 |
| rs10036727 | 0.9596 | 0.1516 | 0.713 | 1.292 | 0.7857 |
| rs1421763 | 1.029 | 0.1671 | 0.7417 | 1.428 | 0.8633 |
| rs9688032 | 1.013 | 0.1511 | 0.7531 | 1.362 | 0.9328 |
| rs11749001 | 0.9943 | 0.1969 | 0.676 | 1.463 | 0.977 |

**Supplementary Table 8.**
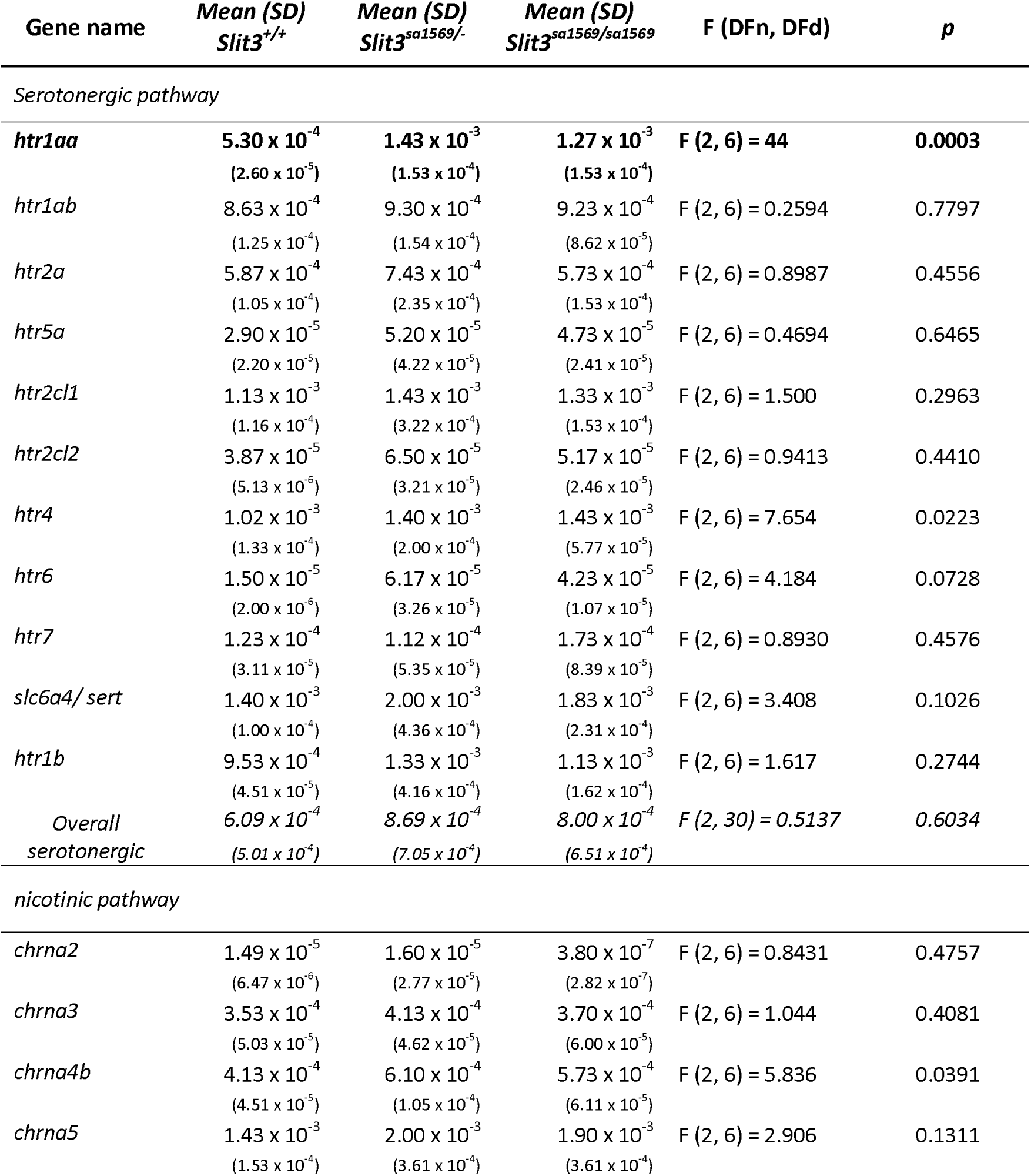
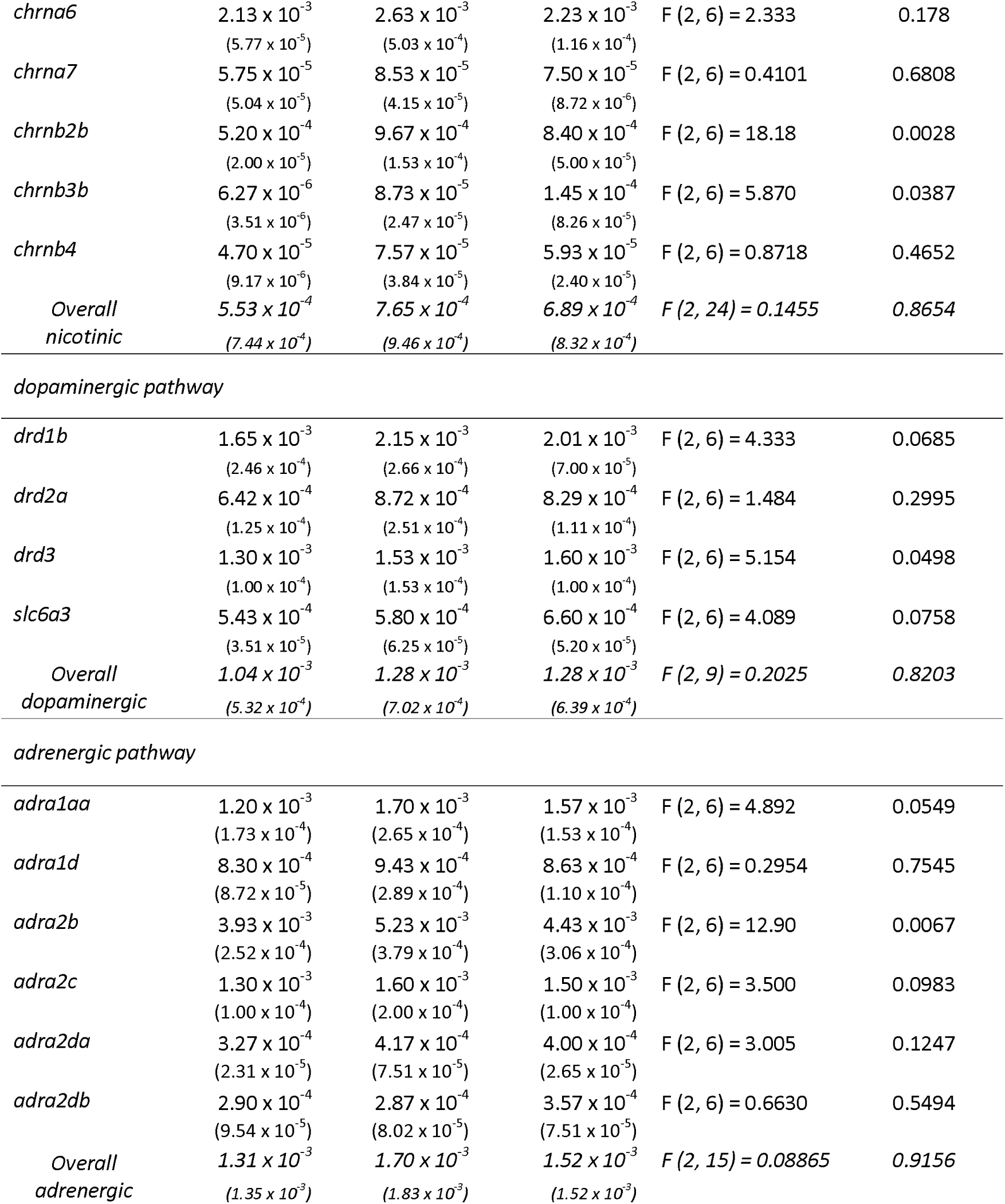
Full results for quantitative real-time PCR analysis of five day old wild type *slit3^sa1569+/+^*, *slit3^sa1569/+^* heterozygous and *slit3^sa1569/sa1569^* homozygous mutant larvae (Total n=30, 3 samples per experimental group with n=10 embryos per sample). Only *htr1aa* ([F(2,6)=44], p=0.0003) showed a significant difference across genotypes after correcting for multiple testing (p < 0.001). DFn and DFd represents degree of freedom for the numerator and denominator of the F ratio.

